# Single-cell analysis of prostaglandin E2-induced decidual cell differentiation: does extracellular 8-Br-cAMP cause artifacts?

**DOI:** 10.1101/2020.09.18.304212

**Authors:** Daniel J. Stadtmauer, Günter P. Wagner

**Affiliations:** Department of Ecology and Evolutionary Biology, Yale University, New Haven, CT, USA; Systems Biology Institute, Yale University, West Haven, CT, USA; Department of Obstetrics, Gynecology and Reproductive Sciences, Yale School of Medicine, New Haven, CT, USA; Department of Obstetrics and Gynecology, Wayne State University, Detroit, MI, USA

**Keywords:** prostaglandin E2, natural ligand, decidua, decidual stromal cell, endometrial stromal fibroblast, PTGER2

## Abstract

Development of the uterine decidua, the transient maternal tissue contacting the fetus during extended gestation, is the hallmark of reproduction in many placental mammals. Differentiation of decidual stromal cells is known to be induced by stimuli that activate the nuclear progesterone receptor and the cyclic AMP/protein kinase A (cAMP/PKA) pathways. The nature of the stimulus upstream of PKA has not been clearly defined, although a number of candidates have been proposed. To bypass this uncertainty for in vitro experiments, direct addition of membrane-permeable cAMP along with progestin has been the prevailing method. Phylogenetic inference suggests that the inflammatory eicosanoid prostaglandin E2 (PGE2) was the stimulus that ancestrally induced decidualization. Accordingly, we developed a protocol to decidualize human endometrial stromal fibroblasts using progestin and PGE2 and analyzed the response in comparison with a cAMP-based protocol. Transcriptomic comparison reveals a common activation of core decidual cell genes between both treatments, and a set of senescence-related genes exaggerated under cAMP treatment. Single-cell transcriptomic analysis of PGE2-mediated decidualization revealed a major transcriptomic transition between an early activated cell state and a differentiated decidual state, but notably did not identify a developmental trajectory representing a distinct senescent decidual state as reported in recent literature. Furthermore, investigation of the signal transduction process underlying PGE2-mediated decidualization showed that it depends upon progestin-dependent induction of PGE2 receptor 2 (PTGER2 aka EP2) and PKA, the kinase activated by PTGER2. This progesterone-dependent induction of PTGER2 is absent in the opossum, a species incapable of decidualization. Together, these findings suggest that the origin of the decidual cell type involved the evolution of progesterone-dependent activation of the PGE2/EP2/PKA axis. We propose the use of PGE2 for in vitro decidualization studies as a potentially more physiological model than 8-Br-cAMP.

## Introduction

The uterine decidua, the transient maternal tissue that forms the maternal side of the fetal- maternal interface, plays an essential role in the reproduction in mammals with invasive placentation (Gellersen and Brosens, 2014). The decidua is a novel tissue which evolved in the stem lineage of placental (eutherian) mammals (Mess and Carter, 2006), and was a key innovation for the evolution of the elaborated placenta of eutherians (Chavan et al. 2017; Stadtmauer and Wagner 2020). Decidual development entails the differentiation of the endometrial stromal fibroblasts (ESF) into mature decidual stromal cells (DSC). The ability of endometrial stromal fibroblasts to differentiate into DSC is an evolutionarily derived trait (Park et al., 2016; Erkenbrack et al., 2019), and the stromal cells that are able to form DSC are called “neo-ESF” (Kin *et al.,* 2016). Neo-ESF and DSC evolved from an ancestral endometrial stromal cell type unable to decidualize called “paleo-ESF.” Paleo-ESF are still found in the opossum, *Monodelphis domestica* (Kin *et al.,* 2016), and likely exist in other marsupials. It follows that comparison of the response of paleo-ESF and neo-ESF to deciduogenic stimuli is a way to identify the molecular key innovations of neo-ESF to enable differentiation into a DSC. Here we investigate which signal, in addition to progesterone (P4), was likely the ancestral trigger for decidual differentiation and come to the conclusion that PGE2 likely is the “natural” ligand for decidualization. We have previously described the response of paleo-ESF to this ligand (Erkenbrack et al., 2018), and here we describe the response of the human ESF to PGE2 and progestin. We propose that one of the unique functions of neo-ESF is the specialized progesterone-dependent response to PGE2 identified in this study.

A large amount of molecular biology has been done on the mechanisms underlying decidualization (Gellersen and Brosens, 2014). A broadly accepted model suggests that decidualization results from the convergence of two signaling pathways, progesterone signaling via isoform A of the nuclear progesterone receptor, PR-A, and the cAMP/PKA pathway (Brosens et al., 1999; Christian et al., 2001). While many other factors have been shown to influence or are associated with decidualization, the main evidence for the two signaling pathway model is that a membrane-permeable form of cAMP (8-Bromo-cAMP) and progesterone are sufficient to decidualize human endometrial stromal fibroblasts in vitro. It is less clear what the ligand is that activates the cAMP/PKA pathway in vivo. Candidates include: gonadotropins such as follicle stimulating hormone (FSH), luteinizing hormone (LH), or human chorionic gonadotropin (hCG) (Tang and Gurpide, 1993), relaxin (RLN) (Telgmann et al., 1997), corticotropin releasing hormone (CRH) (Ferrari et al., 1995) and PGE2 (Frank et al., 1994) (see review in Gellersen and Brosens, 2014). However, the receptors of gonadotropins, RLN, and CRH are either not or only marginally (*RXFP1*) expressed in cultured human ESF, both immortalized as well as primary cells (Wagner, unpublished), and are thus unlikely to be critical for decidualization. Also, other candidates have evolved more recently, for instance hCG only exists in anthropoid apes (Maston and Ruvolo, 2002) and therefore cannot be the ancestral stimulus for decidualization. In contrast, PGE2 receptors are broadly expressed across mammalian ESF, from opossum to human, with the exception of species that secondarily lost invasive placentation, e.g. the cow (Kin et al., 2016). In addition, PGE2 is present at the implantation site after embryo attachment in a wide range of eutherian mammals (Table S1), and even in the opossum (Griffith et al., 2017; Erkenbrack et al., 2018). This means that PGE2 was a signal present in the uterus at the time of embryo attachment even before the evolution of the DSC, and the evolution of decidualization would have entailed evolution of a new response to PGE2, rather than a new decidualizing signal. This follows from the finding that decidual cells evolved after the most recent common ancestor of opossum and placental mammals (Kin et al., 2014), and that embryo attachment in the opossum is also associated with PGE2 production. Hence, we investigated whether PGE2, in addition with P4, is sufficient for in vitro decidualization of human ESF. PGE2 has been implicated in decidualization before (Frank et al., 1994; Brar et al., 1997), and was then interpreted as a modulating factor enhancing decidualization.

The widely-used standard protocol uses extracellular cAMP to activate the PKA pathway could lead to artifacts because the cell is unable to internally regulate the level of cAMP. In fact, here we show that the cAMP-based protocol causes the expression of a large set of genes, including markers of a recently reported senescent decidual cell state (snDSC) (Lucas et al., 2020), which are much more highly expressed compared to our PGE2-based protocol. To investigate this interpretation we perform single-cell transcriptome analysis on PGE2/MPA-stimulated human ESF and find that in fact during in vitro decidualization ESF are producing two kinds of cells, an activated ESF population and mature DSC, where the former has some characteristics of senescent cells but does not seem to represent a distinct population of senescent cells.

## Materials and Methods

### Cell Culture and in Vitro Decidualization

Immortalized human endometrial stromal fibroblasts (equivalent to ATCC line CRL-4003) were cultured in growth media containing the following: 1x antibiotic-antimycotic (15240, Gibco; contains penicillin, streptomycin, and Amphotericin B), 15.56 g/L DMEM/F-12 without phenol red (D2906, Sigma Aldrich), 1.2 g/L sodium bicarbonate, 10 mL/L sodium pyruvate (11360, Thermo Fisher), and 1 mL/L ITS supplement (354350, VWR) in 10% charcoal-stripped fetal bovine serum (100-199, Gemini). Media was replaced every 3-4 days during cell growth.

For in vitro decidualization, cells were grown in a base medium containing antibiotic- antimycotic (15240, Gibco), 15.56 g/L DMEM/F-12 (D8900, Sigma Aldrich), and 1.2 g/L sodium bicarbonate in 2% fetal bovine serum (100-106, Gemini). Additives included medroxyprogesterone acetate at a concentration of 1 μM, prostaglandin E2 at a concentration of 1 μM, or 0.5 mM 8- bromoadenosine 3’-5’-cyclic monophosphate. Unless otherwise noted, media was changed daily during the decidualization process. Decidualization was begun when cells had reached approximately 80% confluency in 6-well (9 cm^2^) culture plates.

For inhibitor experiments, in vitro decidualization was conducted with the addition of either 1 μM PTGER2 inhibitor (PF-04418948, Sigma), 1 μM PTGER4 inhibitor (ER-819762, Sellek), or 5 μM of the PKA inhibitor H-89 (B1427, Sigma). Concentrations were determined based upon published literature (PF-04418948: Birrel et al., 2012; ER-819762: Chen et al., 2010; H-89: Yoshino et al., 2003; Matsuoka et al., 2010) and preliminary experiments.

### Bulk RNA Sequencing

RNA was isolated from cells lysed using Qiagen Buffer RLT and processed using the RNeasy Mini Kit (74104, Qiagen) according to the manufacturer’s specifications. Transcriptome libraries were generated from isolated RNA by the Yale Center for Genomic Analysis using poly-A selection and sequenced using an Illumina NovaSeq.

Sequencing reads were mapped to the Ensembl human transcriptome GRCh38 version 96 using the pseudoalignment software kallisto (version 0.42.5; Bray et al., 2016). Transcript read counts were normalized using the transcripts per million (TPM) metric (Wagner et al., 2012) for comparison between samples. The Pvclust software package (Suzuki et al., 2006) was used to ascertain the uncertainty in hierarchical clustering of samples.

Principal component analysis was conducted on a subset of transcript per million transcriptomes, only taking into consideration genes with an average expression of between 3 and 1000 TPM across all conditions. Genes were subset to only those with false discovery rate < 10^-6^, fold change > 1.5 (treatment/control expression > 1.5 for upregulated or control/treatment > 1.5 for downregulated), and excluding genes designated “off” (TPM < 3) in both conditions.

Differential expression analysis was conducted using edgeR (version 3.28.0; Robinson et al., 2010). The resulting differentially expressed genes were filtered to only those expressed at greater than 2 transcripts per million, an operational threshold for being considered “on” (Wagner et al., 2013).

Furthermore, only genes with a false discovery rate lower than 10^-6^ and an average fold change of greater than 1.5 between treatment and control (treatment/control expression > 1.5 for upregulated, or control/treatment > 1.5 for downregulated) were considered. For analyses that only took protein-coding genes into consideration, filtering using the Ensembl BioMart “protein_coding” gene type annotation reduced the total number of genes from 40275 to 22688, and transcripts per million were recalculated accordingly. The number of genes that met all of the above criteria was 2668.

K-means clustering of genes by square root-transformed TPM values averaged for each treatment condition was conducted to generate a number of clusters ranging from 4 to 10. Following manual comparison, k = 5 clusters appeared to capture the most prominent patterns from our heatmap including all genes, so more extensive analysis was conducted on these five clusters. Pathway analysis and gene ontology annotation were performed using Enrichr software (version 2.1; Kuleshov et al., 2016).

### Single-cell RNA Sequencing

Single-cell RNA sequencing was conducted on endometrial stromal fibroblasts decidualized in vitro for 2 or 6 days by daily administration of media containing 1 μM PGE2 and 1 μM MPA. Single- cell suspensions were prepared by a protocol modified from the 10X Genomics Sample Preparation Demonstrated Protocol for Cultured Cells (CG00054 Rev. B). Briefly, cells were rinsed with TrypLE express (12604, Thermo Fisher) and treated with TrypLE express for 5-10 minutes at 37°C. 10 mL growth medium was added and the cells were pelleted by centrifugation for 5 minutes at 400 g, then resuspended in 5 mL growth medium by pipetting up and down 10-20 times. Next, an 18-gauge needle was used to transfer the cells onto a 40-μm strainer, and the flowthrough was centrifuged and resuspended in PBS containing 0.04% bovine serum albumen (A9647, Sigma). The resulting suspension was passed through a FlowMi 30-μm cell strainer and used for single-cell library preparation using the 10X Chromium platform (3’ chemistry, version 3) and sequenced using an Illumina NovaSeq apparatus. In total, 10073 unique capture bead barcodes (each roughly corresponding to a unique cell) were sequenced, composed of 4706 at 2 days and 5367 at 6 days with an average of 34364 reads per cell.

Initial alignment of raw sequencing reads to the human genome (GRCh38 3.0.0 preparation by 10x Genomics), filtering, barcode and unique molecular indicator counting was conducted using 10X Genomics cellranger software version 3.1.0. Resulting feature-barcode matrices were processed using scprep software version 1.0.5 (github.com/KrishnaswamyLab/scprep). In attempt to remove data points originating from beads that had captured ambient RNA (low library size), multiple cells (high library size), or dying cells having lost membrane integrity (disproportionately high mitochondrial genes), data were further thresholded based on library size (between the 16^th^ and 90^th^ percentile of features/library for 2-day treated samples, and 9^th^ and 90^th^ percentile of features/library for 6-day treated samples), and below the 93^rd^ percentile of mitochondrial gene content of the library after examination of distributions (Figure S3). Genes expressed in fewer than 5 cells from either sample were excluded from the analysis. The result was a total of 7279 cells and 16496 genes considered in downstream analyses.

Read counts were normalized to counts per million and square root transformed. Low-dimensional embedding was performed using PHATE (version 1.0.5; Moon et al., 2019) For the purpose of visualizing individual gene expression, denoising was performed using MAGIC (version 2.0; van Dijk et al., 2018). The statistical dependency between genes or between genes and pseudotemporal scores was quanified using k-nearest neighbor conditional density resampled estimate of mutual information (knn-DREMI) and rescaled for visualization using DREVI where noted (Krishnaswamy et al., 2014; van Dijk et al., 2018).

Cells were partitioned using k-means clustering with increasing k parameters (Figure S4) and the lowest number of clusters was chosen for which each prominent region of the embedding was partitioned into a unique cluster, but before a precipitous drop in silhouette score. Marker genes for each cluster were identified using scprep’s differential expression function which performed Welch’s *t*- test between cells within and without each cluster.

Unbiased annotation of single cells with their most similar bulk RNA-seq treatment condition (base media, PGE2+MPA, or cAMP+MPA) was performed using SingleR (v1.0.6; Aran et al., 2019). Trajectory inference was performed individually on 2- and 6-day treated samples using Slingshot (Street et al., 2018). A starting cluster of most ESF-like cells, as determined by SingleR and visualization of ESF marker genes, was used as a constraint on the Slingshot algorithm.

### Senescence-Associated β-Galactosidase

Human endometrial stromal fibroblasts were cultured in base differentiation media with one of the following sets of additives: none (control), 1 μM palbociclib (PD0332991, Sigma) plus 1 μM PGE2 and 1 μM MPA, 1 μM PGE2 plus 1 μM MPA, 0.5 mM cAMP plus 1 μM MPA, or 1 μM MPA. Media was replenished daily over the course of six days. At the end of the time course, senescence-associated β-galactosidase activity was ascertained using a staining kit (9860S, Cell Signaling) following the manufacturer’s instructions. In this protocol, cells were rinsed in PBS and treated with fixative solution for 10-15 minutes, rinsed with PBS twice, and incubated overnight at 37°C before imaging.

## Results

### Progestin + Prostaglandin E2 Treatment Activates Classical Hallmarks of Decidualization

Immortalized human endometrial stromal fibroblasts were treated with either progestin (MPA, metroxyprogesterone acetate) and prostaglandin E2, progestin and cyclic AMP, or base culture media alone. PGE2+MPA treatment induced a morphological change from an elongate fusiform morphology to a more quasi-epithelial, laterally expanded shape (Figure 1a). They did not become globular to the same extent observed in response to treatment with progestin and cyclic AMP (Figure 1a).

**Figure 1.**
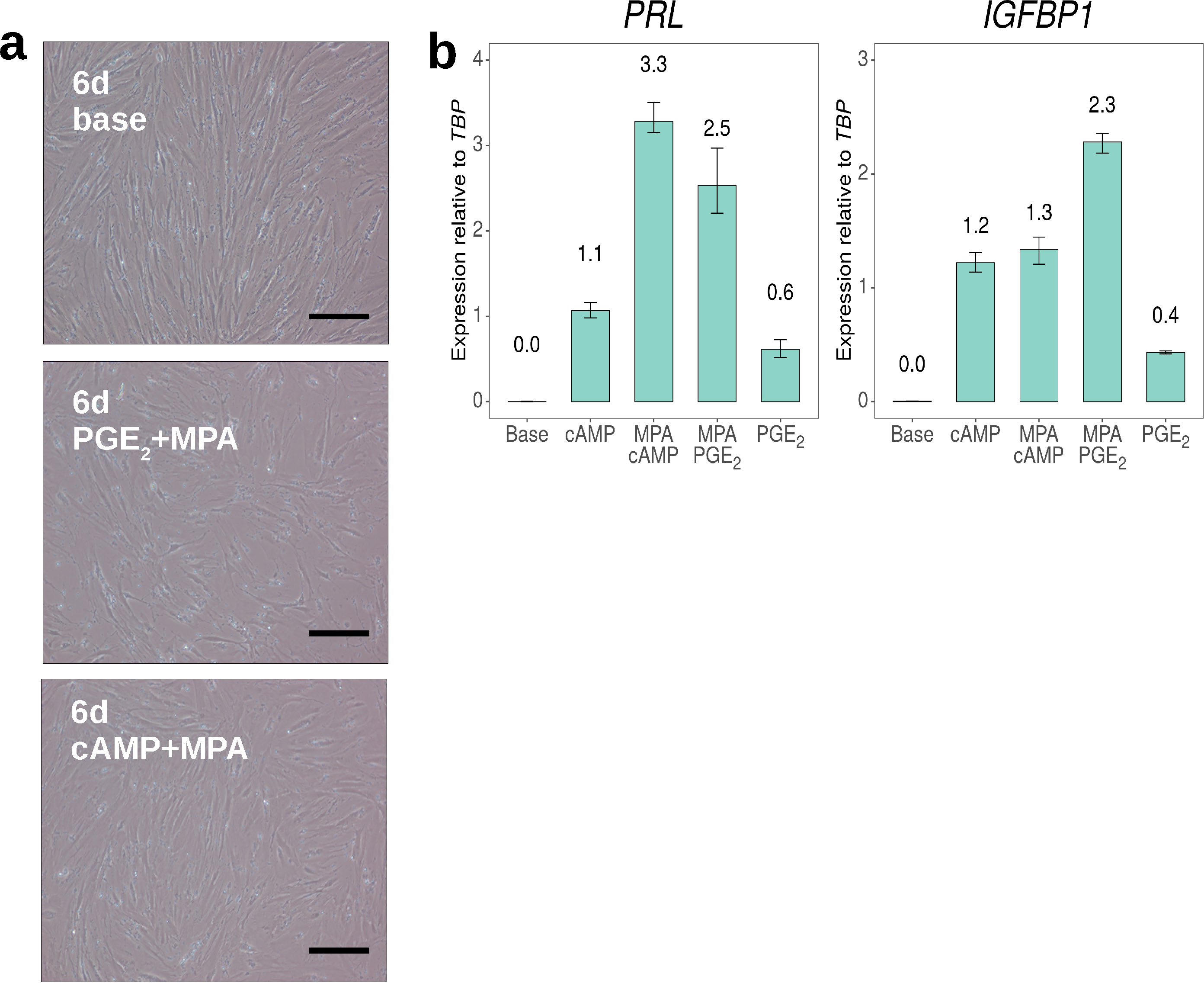
In vitro decidualization of human ESF using PGE2+MPA. **a.** Cells treated with culture media for 6 days (left) retained spindle morphology, whereas cells treated with MPA+PGE2 for 6 days became wider (right). Scale bar = 20 μm. **b.** qPCR quantification of decidual markers PRL and IGFBP1 in response to various decidualization regimes, measured as linearized expression relative to control gene TBP. Experimental n = 2 for cAMP+MPA and PGE2+MPA treatments, experimental n = 1 for all other treatments, and technical n = 3 for all conditions.

Nevertheless, the expression of key marker genes *prolactin* (*PRL*) and *insulin-like growth factor binding protein 1* (*IGFBP1*) were upregulated to a similar extent in response to both treatments as determined by quantitative real-time PCR (qPCR) (Figure 1b).

### PGE2+MPA and cAMP+MPA Treatments Modulate an Overlapping Set of Genes

Bulk transcriptome sequencing was conducted on cells exposed to both decidualization regimes, as well as to a base media control. Hierarchical clustering analysis found with high confidence that PGE2+MPA-treated cells clustered with cAMP+MPA-treated cells, to the exclusion of the untreated control group (Figure S1). Principal component analysis (Figure 2a) was used to visualize the collective similarity of the three gene expression states. Transcriptomes were found to separate along the first two principal component axes primarily by treatment condition. The first axis, explaining 47.3% of the variance, separated the two decidualization treatments from the control treatment most notably. The second principal component axis, explaining 26.6% of the variance, primarily separated PGE2+MPA treatment from the cAMP+MPA samples. This pattern suggests that cells from both treatments differ in similar ways from ESF and also display expression differences specific to their treatment.

**Figure 2.**
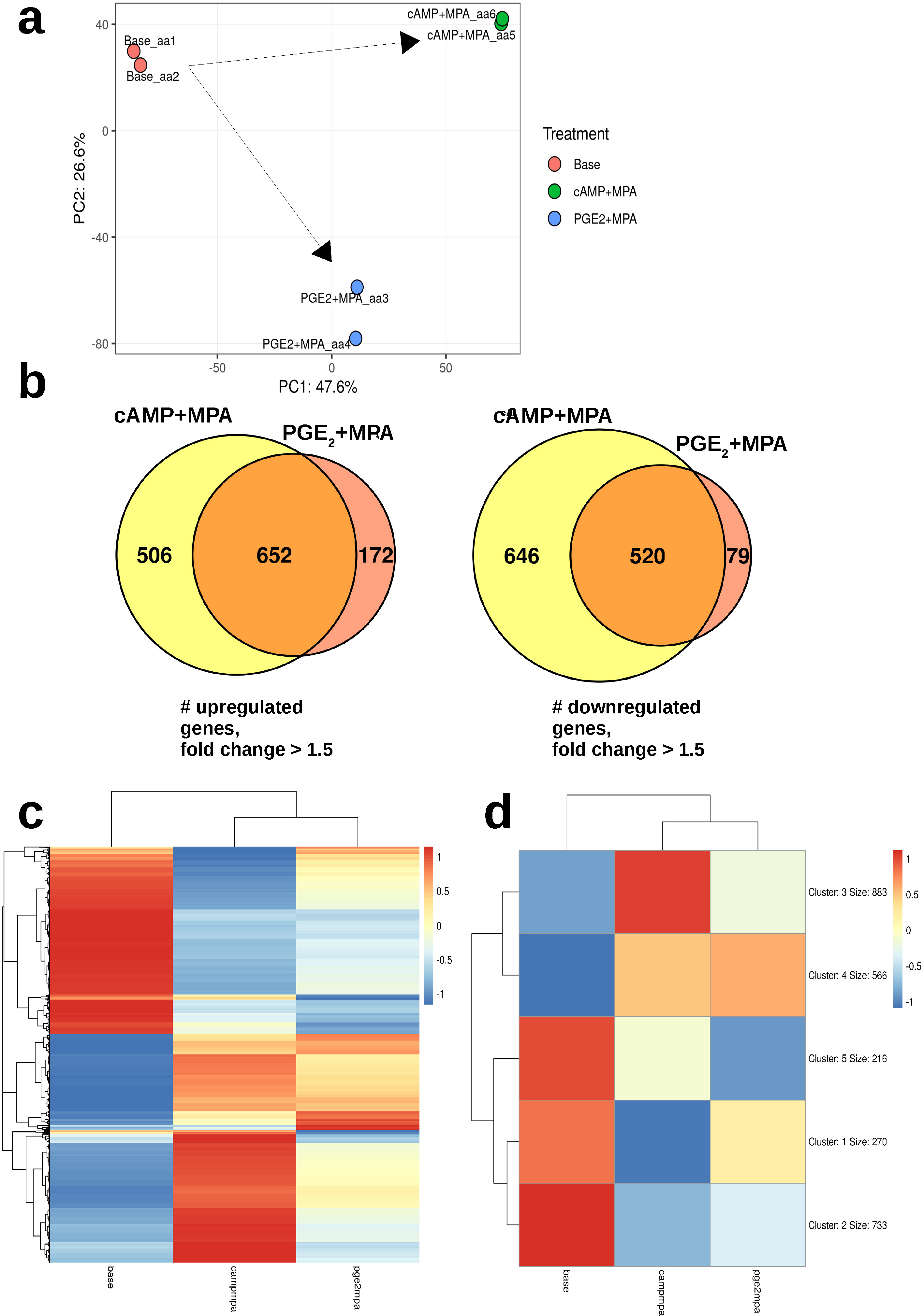
Bulk RNA sequencing analysis. **a.** Principal component analysis conducted on protein-coding genes. Transcriptomes cluster by treatment, with PC1 representing a putative decidual differentiation axis. **b.** Venn diagrams of genes differentially upregulated or downregulated in response to cAMP+MPA or PGE2+MPA treatments with respect to control treatment. **c.** Heatmap of differentially expressed protein-coding genes (n = 2 per condition). **d.** K- means clustered (k = 5) heatmap of differentially expressed protein-coding genes showing 5 major patterns of comparative gene expression among the 3 treatment conditions (n = 2).

Differential gene expression analysis was conducted on all pairwise combinations of the three treatment regimes: PGE2+MPA, cAMP+MPA and control. There was substantial overlap of 652 genes found to be upregulated under cAMP+MPA treatment (56% of 1158 genes) as well as under PGE2+MPA treatment (79% of 824 genes) (Figure 2b). The overlap of downregulated genes (520 of 1166, 45%, under cAMP+MPA, 520 of 599, 87%, under PGE2+MPA) was similarly large. In both cases the set of genes differentially expressed in response to cAMP+MPA treatment was larger than the number of genes uniquely responsive to PGE2+MPA (506 versus 172 genes uniquely upregulated, 646 versus 79 genes uniquely downregulated). This pattern suggests that cAMP is inducing expression of genes that are not affected by PGE2. The question, addressed below, is whether that means that PGE2 is only inducing a subset of decidualization-related genes, or whether cAMP is causing artifactual gene expression.

### Both PGE2 and cAMP Activate the Decidual Cell Core Regulatory Network

Several prominent patterns emerged in the gene expression of cells under both decidualizing treatments. If expression scores are divided into “low/off” “medium” and “high” expression categories (Table S2), there are 27 possible patterns among the three treatments, 24 of which are differential (excluding identical categories in all 3 groups). Heatmap clustering of differentially expressed genes (Figure 2c) revealed, however, that some of these conceivable combinations were much more prominent in the data than others. To investigate this further, k-means clustering of genes was conducted using a range of hyperparameters from k = 3 to k = 10 clusters, and it was concluded that the data were best represented by five major patterns of differential gene expression between the three treatment conditions (Figure 2d). The full output from this analysis is provided as Supplemental Data.

Two of the resulting gene clusters represented genes lowly expressed in the control treated group, and three represented genes highly expressed in ESF but downregulated in response to either cAMP+MPA or PGE2+MPA treatment. Of the former two, Cluster 4 included 566 genes that generally were moderately to highly upregulated in response to both decidualizing regimes. This cluster included almost all of the genes that have been identified as belonging to the decidual cell type’s core gene regulatory network (Erkenbrack *et al.,* 2018; Liang, 2018), including transcription factors c-Fos *FOS*, nuclear receptor related 1 protein *NR4A2*, forkhead box protein O1 *FOXO1*, odd-skipped-related 2 *OSR2*, hypoxia-inducible factor 1-alpha *HIF1A*, zinc finger and BTB domain-containing protein 16 *ZBTB16*, heart- and neural crest derivatives-expressed protein 2 *HAND2*, Krueppel-like factor 9 *KLF9*, zinc finger protein 331 *ZNF331*, and zinc fingers and homeoboxes protein 2 *ZHX2.* This cluster also included phenotypic effector genes encoding signaling peptides associated with decidual cells such as *PRL*, *IGFBP1*, left-right determination factor 2 *LEFTY2*, bone morphogenetic protein 2 *BMP2*, and somatostatin *SST* (Table S3). This consistent pattern suggested that a true differentiation event to the decidual cell type had taken place in both treatment regimes.

The latter three gene clusters consisted of genes highly expressed in ESF and more lowly expressed under either treatment. Of these, Cluster 2 included 733 genes that were similarly downregulated by both decidualization regimes. Pathway analysis of this cluster returned terms primarily associated with cytoskeletally-mediated fibroblast functions, such as “focal adhesion” (KEGG, p = 8.0×10^-12^), “tight junction” (KEGG, p = 2.2×10^-4^), and “regulation of actin cytoskeleton” (KEGG, p = 2.0×10^-6^). These represented genes mediating cellular interactions including thrombospondin-2 *THBS2*, tubulins *TUBB6*, *TUBB4B*, *TUBA1C*, and *TUBA1B*, ezrin *EZR*, and the claudin *TMEM47*. Furthermore, genes related to cell proliferation, including terms related to “cell cycle” (KEGG, p = 4.3×10^-5^), including mitosis (GO:0000070; GO:0044772; 0000086; 0044839) and cytokinesis (GO:0000281; GO:0061640) were included in this cluster, indicative of the cell cycle exit expected to accompany a differentiation event (Takano et al., 2007).

There were two patterns of genes highly expressed in endometrial stromal fibroblasts and downregulated in one or the other decidualization regime. One of these, Cluster 5, consisted of 216 genes for which the PGE2+MPA treatment led to stronger downregulation than cAMP+MPA. This included growth factors such as pleiotropin *PTN* and neuregulin 1 *NRG1,* and the GDNF receptor *GFRA1* (Table S5). This pattern also applied to genes involved in extracellular matrix production associated with decidualization such as osteopontin *SPP1* and hyaluronic acid synthase *HAS2* (Table S5).

Lastly, Cluster 1 comprised 270 genes highly expressed in ESF that were more strongly downregulated by cAMP+MPA treatment than PGE2+MPA treatment. Some of the genes most specifically affected by cAMP+MPA (i.e. non-significant difference between PGE2+MPA and base treatments) were genes having to do with translation such as the aminoacyl-tRNA synthetases *WARS*, *YARS*, *MARS, GARS*, and *SARS.* This cluster of also included the matrix proteoglycans testican *SPOCK1* and versican *VCAN,* as well as the mesenchymal regulator SLUG *SNAI2*. This cluster also contained genes belonging to the KEGG pathway “PI3K-Akt signaling pathway” (p = 9.3×10^-4^), including the GTPase *HRAS*, PH domain and leucine-rich repeat protein phosphatase *PHLPP1*, and fibronectin *FN1*.

### Genes Specifically Induced by cAMP+MPA

Genes whose expression was specifically induced by cAMP+MPA (Figure 2b) fell into Cluster 3 under k-means clustering (Figure 2d). Notably included in this cluster of 883 genes were almost all of the markers identified in a previous study (Lucas et al., 2020), and were proposed to represent a senescent subpopulation of decidual cells. These included ABI family member 3 binding protein *ABI3BP*, A disintegrin and metalloproteinase with thrombospondin motifs 5 *ADAMTS5*, cell migration- inducing and hyaluronan-binding protein *CEMIP* (*KIAA1199*), clusterin *CLU*, type II iodothyronine deiodinase *DIO2*, insulin-like growth factor 2 *IGF2*, lumican *LUM*, epididymal secretory protein E1 *NPC2*, and the SRY-related HMG-box transcription factor *SOX4* (Table S4). Lucas and colleagues (2020) have proposed that decidualization leads to two fates, decidualization and senescence, with both playing important biological roles in implantation. The preferential elevation of this signature in cAMP+MPA decidualization prompted further investigation into whether such a senescent cell population was absent in PGE2+MPA-decidualized cells.

As a first test for cellular senescence, chromogenic staining for senescence-associated β- galactosidase (SaβG) activity was conducted on endometrial stromal fibroblasts decidualized for 3 or 6 days by cAMP+MPA or PGE2+MPA regimes (Figure S2). Highest β-galactosidase activity was observed in the cells treated for 6 days with decidual stimulus, regardless of whether it was PGE2+MPA or cAMP+MPA. On the other hand, cells treated with MPA alone for an equivalent duration showed fewer SAβG^+^ cells. Cells decidualized with PGE2+MPA in the presence of palbociclib, an inhibitor of CDK4/CDK6 previously used to increase susceptibility of cells to senescence (Brighton et al., 2017), showed no noticeable effect in our semi-quantitative test. We interpret this result as indicating that under both treatment regimes the cells experience cellular stress, probably caused by the energy demand associated with transcriptional reprogramming.

### Single-Cell Transcriptomic Analysis of Prostaglandin E2-Induced Decidualization

Since gene expression associated with senescent DSC was largely limited to cAMP+MPA treatment in our bulk RNA-seq experiments but simple SAβG tests were inconclusive with respect to the presence of senescent cells, we investigated the question of whether PGE2+MPA treatment leads to the formation of a senescent cell population. We approached this question by single-cell RNA sequencing.

Cell cultures were subjected to single-cell transcriptomic analysis after 2 or 6 days of PGE2+MPA treatment. This design was chosen because according to Lucas et al. (2020), the senescent cells arise after four days of decidualization treatment. Single-cell transcriptomes were embedded in two dimensions, and kernel density estimation, a method to visualize data point density when normally points would overlap in the ordination plot, was used as a first-pass attempt to judge the number of potential cell types by looking for hotspots of cell density (Moon et al., 2018). The prediction was that this number would increase by one after 6 days indicating the emergence of a senescent population.

However, this visualization instead revealed two centers of data density at 2 days, connected by a continuous line, and a single center in the 6-day treated samples (Figure 3a). This is more indicative of a convergence than a bifurcation.

**Figure 3.**
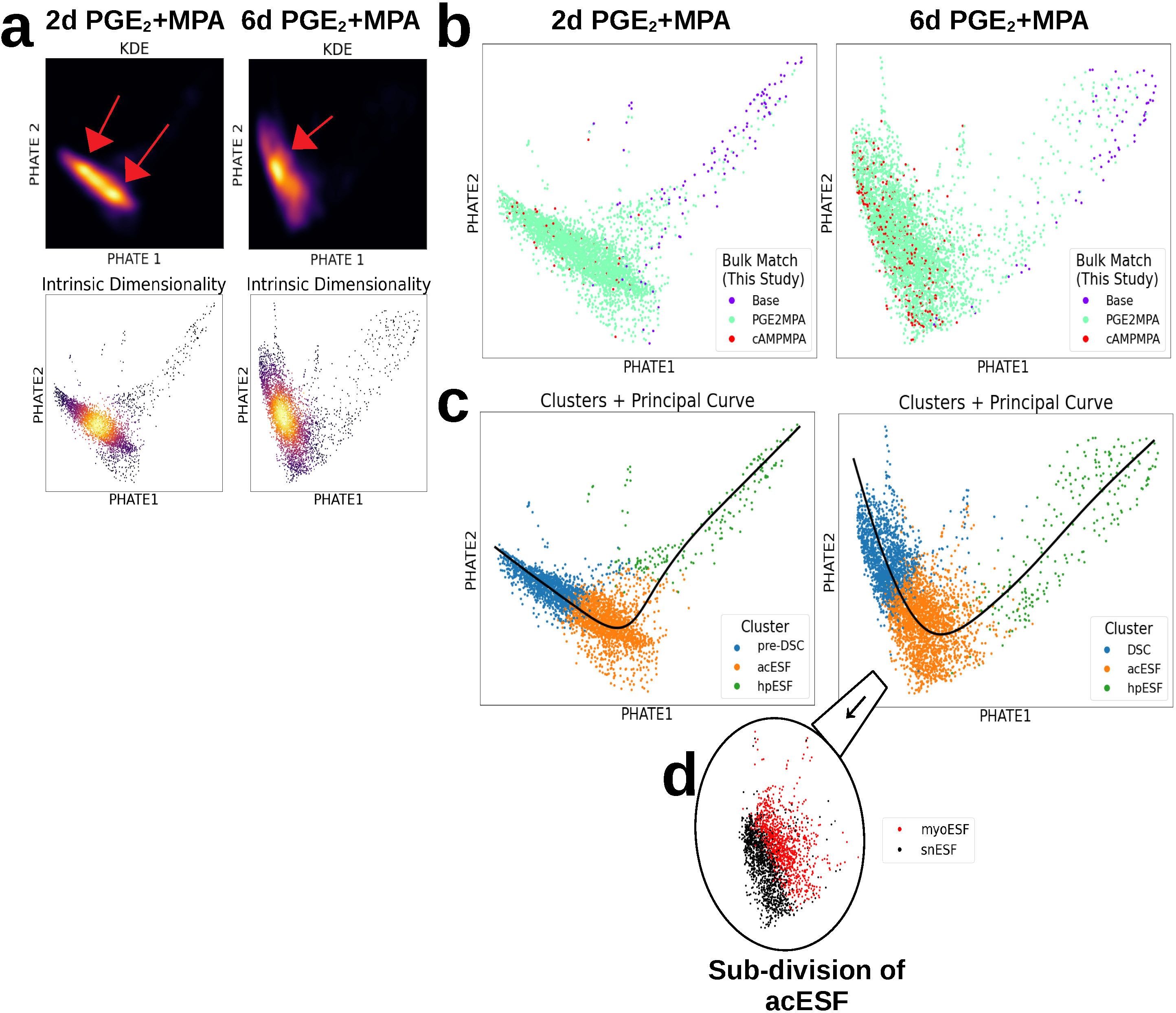
Single-cell RNA sequencing analysis. **a.** Kernel density estimation shows the concentration of most cell density in a region to the lower left of 2-dimensional PHATE embedding, with either two centers of density in the 2-day treated cells (top), or one in the 6-day treated cells (bottom). **b.** All cells from 2-day (left) or 6-day (right) treatments were plotted in a 2-dimensional space and color-coded by best match to bulk RNA-seq treatment conditions (this study). This revealed a curve representing decidual progression from ESF-like (base treatment) to DSC- like (mature) **c.** PHATE visualization for 2- and 6-day PGE2+MPA decidualized cells, paritioned into clusters and overlaid with inferred developmental trajectories. **d.** Subdivision of activated ESF cluster into two partitions, SOX4^+^ snESF and ACTA2^+^ myoESF.

The second preliminary assessment was exploration of intrinsic dimensionality of the data, in order to identify potential branch points: the prediction was that cells in transition between cell types would have more complex transcriptomes (higher intrinsic dimensionality), whereas metastable, differentiated states would have lower intrinsic dimensionality (Moon et al., 2019). This analysis identified a single locus of cells ranked highest in intrinsic dimensionality between the two cell density hotspots at 2 days, and slightly below the major hotspot identified at 6 days (Figure 3a). This indicated that the major transcriptomic transition was occurring in this region.

The next step was the identification of the cell states observed with stages of ESF differentiation. Visualization of cell transcriptomes revealed that the majority of cells lie on or around the identified density hotspots, but that at both time points a sparse string of cells extended out to form a curved tail continuous with the line between the two hotspots. The curve formed by this sparse tail leading into the denser cell area was hypothesized to represent progression along the decidualization pathway. This was tested by assignment of each cell to its transcriptionally most similar group from the bulk RNA-seq experiment, which revealed that the less densely populated end of the curve was indeed most ESF-like (Figure 3b). The more densely-populated end of the ordination plot was most similar to PGE2+MPA-decidualized bulk transcriptomes, with some cAMP+MPA interspersed but not characterizing a distinct region.

Cells were partitioned into transcriptomically similar clusters using k-means clustering (Figure 3c). The optimal number of clusters was determined from a range of 2 to 7 (Figure S4). The first partition recognized was between cells above and below the previously identified point of highest intrinsic dimensionality, which presumably represented the transition point between these two distinct cell states or even cell types. Next, at both time points, a second division occurred at k > 3 clusters between the sparse ESF-like tail and the rest of the accumulation of cells below the transition point. Further divisions sharply reduced clustering silhouette scores, so a total of 3 clusters seems to be the most meaningful classification of cells for both, the 2- and 6-day time points.

Gene expression analysis was used to identify marker genes for each cluster (Supplemental Data) and explore expression of known markers of various DSC and ESF cell states (Figure 4). At the sparsely-populated ESF-like end of the curve, ESF markers such as *VIM*, *EZR* and *TUBB6* (Figure 4a) were most highly expressed at both 2 and 6 days. Furthermore, these cells also had a distinct cell cycle signature (e.g. GO:1901990, “regulation of mitotic cell cycle phase transition”, 2d p = 5.84×10^-15^, 6d p

**Figure 4.**
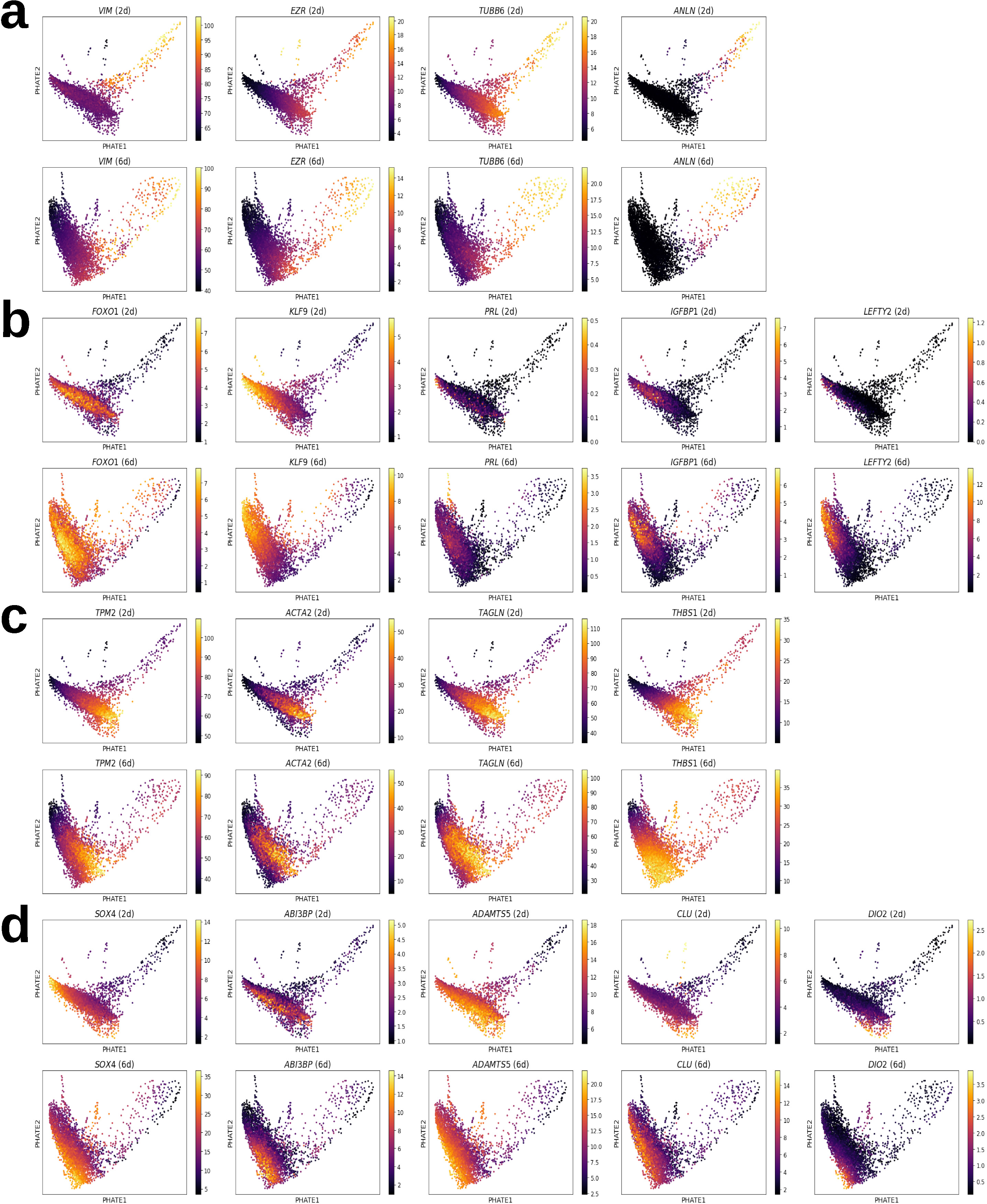
Expression of select genes (MAGIC-denoised square root counts per million) plotted over reduced-dimensionality single-cell data for either 6-day or 2-day time points. **a.** Proliferative ESF marker genes: *VIM*, *EZR*, *TUBB6*, *ANLN*. **b.** Canonical DSC markers: *FOXO1*, *KLF9*, *PRL*, *IGFBP1*, *LEFTY2*. **c.** Pre-decidual ESF marker genes: *TPM2*, *ACTA2*, *TAGLN*, *THBS1* **d.** Markers of proposed progesterone-resistant/senescent/SOX4^high^ DSC: *SOX4*, *ABI3BP*, *ADAMTS5*, *CLU*, *DIO2*.

= 3.79×10^-13^). Indeed, cell cycle scoring identified this tail as cells in S and G2/M phase, whereas the bulk of cells were G1 (Figure S5a). This pattern included the cytokinesis gene anillin *ANLN* (Figure 4a), which was previously identified as a marker of proliferative ESF (Lucas et al., 2018), indicating a distinction between proliferation (highly proliferative ESF; hpESF) and differentiation. Nevertheless, regression out of cell cycle-associated genes (Tirosh et al., 2016), a step sometimes included in scRNA-seq analyses although not always advisable due to its potential to mask biological signal (Luecken and Theis, 2019), failed to remove this tail of ESF-like cells (Figure S5b). This confirmed that the cluster was not a cell-cycle artifact but had a broader and persistent ESF-like gene expression signature.

On the other end of the putative decidualization curve, the cluster of cells above the transition point at 6 days showed expression of decidual genes. This cluster was characterized by the gene set “BMP2-WNT4-FOXO1 Pathway in Human Primary Endometrial Stromal Cell Differentiation” (WikiPathways WP3876, 6d p = 8.03×10^-6^), indicating that these were indeed decidual stromal cells. Decidual cell signaling peptides such as *PRL*, *IGFBP1*, and *LEFTY2* were expressed in this region, and decidual transcription factors appearing earlier such as *FOXO1* and *HAND2* (Supplemental Data) or later such as *KLF9* (Figure 4b). Decidual markers were lowly expressed but detectable at 2 days, and at a much greater magnitude at 6 days. At 2 days, this cluster was predominately consisted of ribosomal genes (e.g. GO:0006412 “translation”, p = 5.85×10^-73^), which was interpreted as a signal of cells still undergoing the reprogramming associated with differentiation. This end of the decidualization curve was also characterized by expression of the prostaglandin E2 receptor 2 *PTGER2*, which was unexpressed in the more ESF-like cells.

The dense cluster of cells located below the point of highest dimensionality, more densely populated at 2 days, were also ESF-like, expressing some ESF markers including tropomyosins *TPM1* and *TPM2,* as well as alpha smooth muscle actin *ACTA2* and transgelin *TAGLN* (Figure 4c) but not the decidual markers *PRL* or *IGFBP1* (Figure 4b). At both 2 and 6 days, this cluster was enriched in gene sets such as “regulation of cell migration” (GO:0030334, 2d p = 1.22×10^-5^, 6d p = 2.19×10^-7^), “homotypic cell-cell adhesion” (GO:0034109, 2d and 6d p = 3.87×10^-6^), and “muscle contraction” (GO:0006936, 2d p = 9.64×10^-11^, 6d p = 1.33×10^-7^). This was interpreted as a pre-decidual ESF state, as it lies between the most ESF-like cells and the point of highest intrinsic dimensionality believed to resemble the transition to later-stage decidualization. To differentiate these early-stage decidualizing ESF from proliferative ESF, we term them activated ESF (acESF).

Finally, marker genes of the senescent/progesterone-resistant decidual cell state identified by Lucas et al. (2020) were examined to see whether their expression characterized a subset of cells arising from our experiments with PGE2. These genes, including *SOX4*, *ABI3BP*, *ADAMTS5*, *CLU* and *DIO2* (see Table S4) were expressed in some cells, and found on day 6 to have a stronger correlated expression with each other than on day 2. Particularly at 2 days, expression of most of these genes spread across the putative branch point rather than being exclusive to one region (Figure 4d), but at 6 days expression of several of these genes was concentrated towards one corner of the acESF cells.

Strongest expression of the fibroblast markers *TPM2*, *ACTA2*, and *TAGLN* was, by contrast, more biased towards the opposite side of this cluster, and *THBS1* was more broadly expressed. Sub-partitioning of the acESF cluster in two divided the cells generally between these two patterns (Figure 3d), and revealed the *SOX4^high^* cells expressed gene sets enriched for “p53 signaling pathway” (KEGG p = 1.80×10^-5^) and “PI3K-Akt signaling pathway” (KEGG p = 6.98×10^-3^). On the other hand, *ACTA2^high^ TAGLN^high^* cells were the source of the highest signal for “muscle contraction” (GO:0006936, p = 5.01×10^-8^), “homotypic cell-cell adhesion” (GO:0034109, p =1.48×10^-12^) and “wound healing, spreading of cells” (GO:0044319, p = 1.46×10^-3^). Due to the large decline in clustering optimality (silhouette score) when calculating k = 4 rather than 3 clusters at 6 days, the distinction between the *SOX4*^high^ and *ACTA2^high^ TAGLN^high^* activated ESF was interpreted as gradation within a cluster rather than discrete clusters.

Finally, having identified three major gene expression patterns by marker gene exploration and comparison to bulk RNA-seq, developmental trajectories were inferred between the three shapes and determine whether a new subpopulation was arising the later time point. Trajectory inference was conducted at both time points, with the constraint that all trajectories must begin at the most ESF-like cell cluster. This resulted in only one principle curve (putative trajectory) in the 2-day PGE2+MPA decidualization sample, and one in the 6-day PGE2+MPA decidualization sample (Figure 3c), i.e. no bifurcation event was predicted using this method. The identified trajectory extended to the most DSC cell cluster and represented the axis of decidualization. Partitioning of the 6-day sample into 4 clusters rather than 3 (one representing snESF) did not affect this outcome.

Our single-cell analysis does not support the model that PGE2+MPA treatment leads to the emergence of a separate decidual cell population after more than 4 days of treatment. If such a population exists, it should be seen in our six day treatment. However, we did pick up a weak signal of ESF, not DSC, expressing the snDSC-associated genes identified by Lucas and colleagues (2020), and enriched for genes in the PI3K/AKT pathway. This raises the possibility that the prominent presence of senescent decidual cells after cAMP+MPA treatment may be an artifact of the use of extracellular cAMP rather than a natural ligand to induce decidualization.

### PGE2-Mediated Decidualization Depends Upon PGE2 Receptor 2 and Protein Kinase A

A series of experiments was conducted to determine the signal transduction pathway by which prostaglandin E2 signaling led to decidualization. Of the two PGE2 receptors known to be expressed in vitro in ESF and DSC, *PTGER2* (a.k.a. EP2) was found to increase over pseudotemporal progression along the axis of decidualization whereas *PTGER4* (a.k.a. EP4) decreased (Figure 5a). Likewise, *PTGER2* expression was positively correlated with *PRL* expression as a proxy for decidualization, whereas *PTGER4* was low in almost all *PRL^+^* cells (Figure 5a). These findings suggested that *PTGER2* activation, and possibly *PTGER4* deactivation, has roles in PGE2-mediated decidualization.

**Figure 5.**
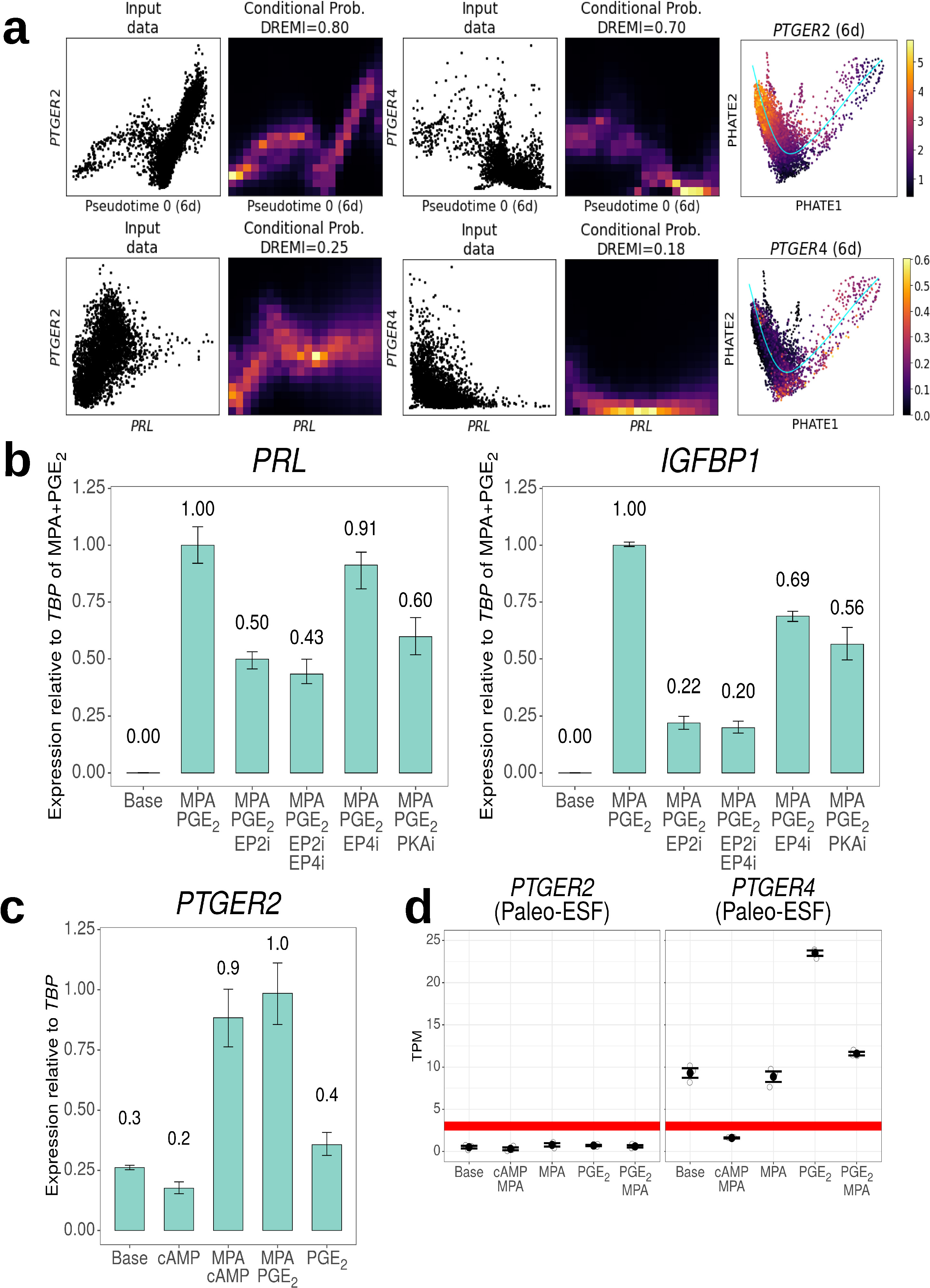
PTGER2 expression is upregulated by progestin and contributes to PGE2- induced decidualization. **a.** Expression of *PTGER2* (left) and *PTGER4* (right) relative to pseudotemporal progression towards DSC or to *PRL* expression in single-cell data. Conditional probability is a density-resampled estimate of mutual information of PTGER2/PTGER4 and the x-axis value. **b.** qPCR quantification of decidual markers in response to PGE2+MPA decidualization as well as inhibitors of PTGER2 (EP2i) or PTGER4 (EP4i), or protein kinase A (PKAi), measured as fold change relative to control gene *TBP* relative to PGE2+MPA group. Experimental n = 2, technical n = 2 for all. **c.** qPCR quantification of *PTGER2* showed upregulation only in the presence of MPA. Experimental n = 2 for both treatments containing MPA, n = 1 otherwise; technical n = 3. **d.** Expression of PGE2 receptors 2 and 4 in *Monodelphis domestica* paleo-ESF (data from Erkenbrack et al., 2018).

To investigate this further, in vitro PGE2+MPA decidualization was conducted in the presence of small molecule prostaglandin E2 receptor antagonists, the PTGER2 antagonist PF-04418948 and the PTGER4 antagonist ER-819762. H-89, an inhibitor of protein kinase A, downstream of PTGER2 signaling, was also administered. Treatment with PTGER2 inhibitor significantly reduced decidual marker genes after 6 days: *PRL* was reduced approximately 50%, and *IGFBP1* approximately 78% relative to uninhibited PGE2+MPA controls (Figure 5b). This effect was marginally improved by treatment with both PTGER2 and PTGER4 inhibitors (57% and 80% reduction, respectively). In contrast, treatment with PTGER4 inhibitor alone resulted in a non-significant 9% reduction in *PRL* expression, and a 31% reduction in *IGFBP1* expression. Administration of the PKA inhibitor H-89 significantly reduced *PRL* expression by 40%, and *IGFBP1* expression by 44%. From these results we conclude that the decidualizing effect of PGE2 is mediated through the PTGER2-PKA axis.

### PGE2 Receptor 2 Is Under Progestin Regulatory Control

Despite the demonstrated importance of PTGER2 to PGE2-mediated decidualization, bulk transcriptomic quantification revealed *PTGER2* to be below or marginally equal to functional levels of expression in endometrial stromal fibroblasts (Figure 5c). However, the gene was clearly turned on after progestin treatment. Expression of *PTGER2* was approximately quadrupled after treatment with MPA plus either PGE2 or cAMP, but not after treatment with either PGE2 or cAMP alone (Figure 5c). This was confirmed in bulk RNA sequencing, which showed an approximate quadrupling of *PTGER2* from ∼3 TPM after control treatment to ∼12 TPM after treatment with MPA and either PGE2 or cAMP. That the effect was indeed due to MPA was confirmed by treatment of ESF for 6 days with MPA alone, after which the expression of *PTGER2* increased 7.14-fold, passing from “off” to “on” (Figure S6).

This effect of progestin on *PTGER2* appears to not be shared by the paleo-ESF cell type.

Analysis of previously published transcriptomes from opossum paleo-ESF treated with MPA alone, MPA+PGE2, or MPA+cAMP (Erkenbrack et al., 2018) revealed that *PTGER2* expression was unaffected by the presence of progestin, and in all cases remained below the operational threshold for being considered “on” (Figure 5d).

## Discussion

In this study we have investigated the biology of PGE2+MPA-induced in vitro decidualization and compared the results with the effects of cAMP+MPA treatment. We found that in vitro PGE2+MPA treatment alone is sufficient to cause the hallmarks of decidualization. Mechanistically, the results suggest that PGE2 mediated decidualization depends on the induction of PTGER2 by progesterone, and proceeds via the EP2-cAMP-PKA axis, while the dominant PGE2 receptor in ESF, EP4, is downregulated.

### Prostaglandin E2 as a Natural Ligand for Decidualization

Whereas previous studies of PGE2-mediated decidualization only performed marker gene and morphological assessment (Frank et al., 1994; Brar et al., 1997), the whole-transcriptome approach paints a more complete picture. The transcriptomic response of endometrial stromal fibroblasts to PGE2+MPA comprised of the widespread activation of genes in the core regulatory network of decidual stromal cells (Table S3). This suggests that the response to PGE2 was a true transformation into the DSC cell type. This conclusion is also supported by the results of our single-cell transcriptomic analysis, which identified a hotspot of cells with high-complexity transcriptomes separating apparently mature DSC from more fibroblast like cells (Figure 3a).

The sufficiency of this molecule, plus the cross-species conservation of prostaglandin signaling in reproduction (Table S1), suggest that both, progesterone and PGE2, can be considered “natural signals” for decidualization. We consider a natural ligand for the differentiation of a cell type to be a shared derived signal that was present and effective at the evolutionary origin of that cell type, and which continues to have an active physiological role in derived species. In decidualization, cAMP mediates a key signal determining decidual cell differentiation but is not the natural ligand itself, whereas PGE2 and progesterone fit the definition of a natural ligand. It has yet to be seen whether other signals, such as those from the local tissue microenvironment including the uterine glands or lumen, also fit this definition. Given its simplicity and the conservation of PGE2 expression in pregnancy among placental mammals, it should be easily applicable to other species.

### PGE2 Signaling through PTGER2 versus PTGER4

Secreted prostaglandin E2 is bound by G-protein coupled receptors in the membrane of the receiving cell. Among the eight members of the prostanoid receptor subfamily, four paralogous receptors are specific for PGE2: PTGER1, PTGER2, PTGER3, and PTGER4 (Breyer et al., 2001). In a resting endometrial stromal fibroblast in culture, only one of these, *PTGER4*, is expressed above the operational threshold for a gene to be considered actively transcribed (Rytkönen et al., 2018). We find that another prostaglandin E2 receptor, *PTGER2*, becomes expressed in human ESF after exposure to progestin.

PTGER2 appears to be functionally adapted to receive sustained PGE2 stimulus, whereas the PTGER4 response is transient and self-limiting (Desai et al., 2000). PTGER4 undergoes more rapid desensitization to ligand binding (Nishigaki et al., 1996), on the order of minutes, due to the action of an additional C-terminal protein domain absent in PTGER2 (Bastepe and Ashby, 1997). Therefore, it is likely that the sequence of events is as follows (Figure 6): At first, PTGER4 is expressed and PTGER2 is absent. Progesterone produced by the corpus luteum after ovulation induces PTGER2. Now, in response to PGE2 stimulus at the time of implantation, PTGER4 becomes quickly desensitized, and signaling through PTGER2 becomes dominant. Indeed, in our bulk RNA-seq data *PTGER4* expression was reduced in response to PGE2+MPA or cAMP+MPA treatment from an average of 5 TPM to 2 or 1 TPM, respectively. Single-cell data also supported this sequence of events, where *PTGER2* was expressed in the mature decidual end of the curve and by contrast *PTGER4* was unexpressed in *PRL*^+^ cells.

**Figure 6.**
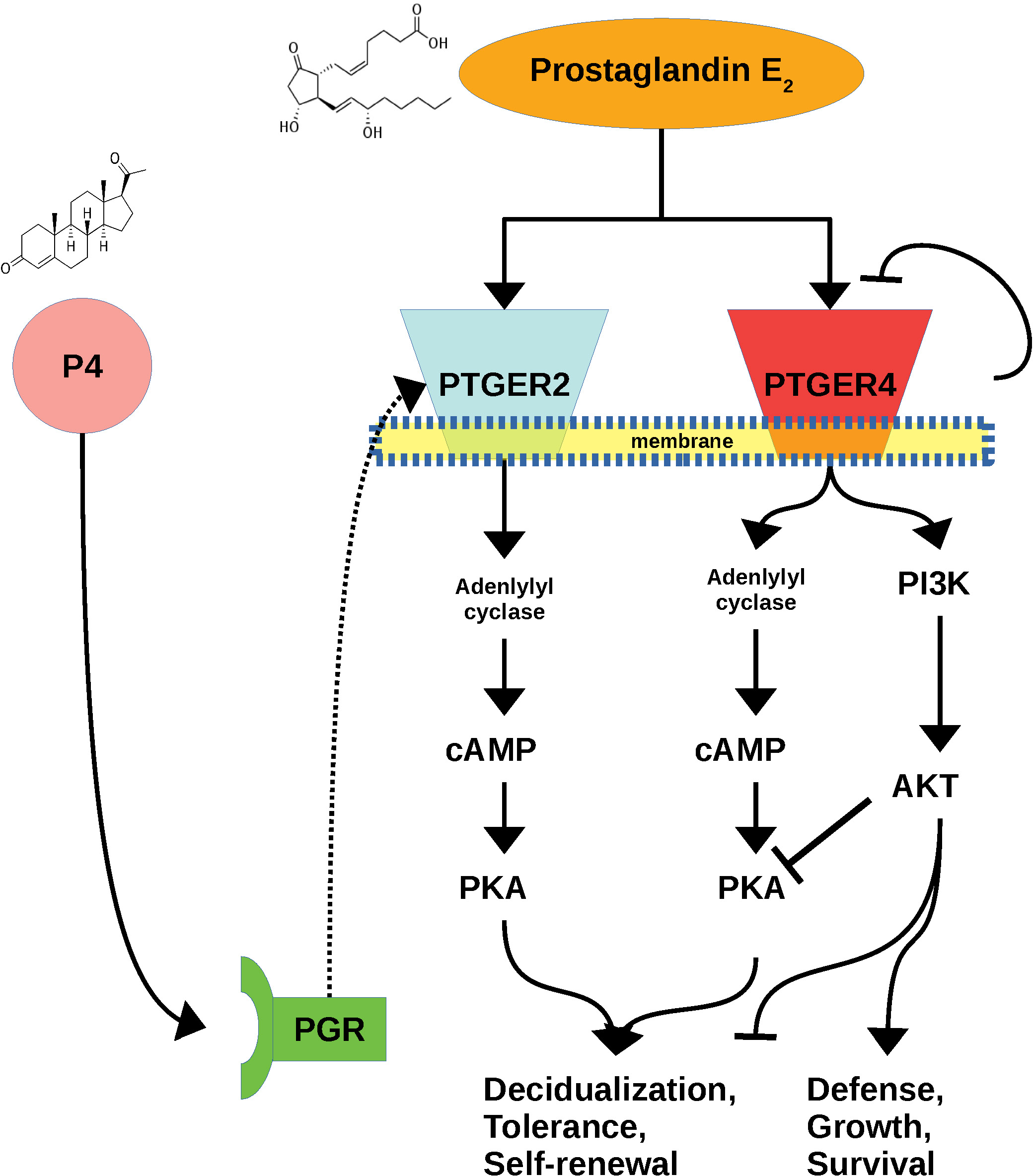
Schematic diagram of induction of decidual cell differentiation. Self-limitation of PTGER4, plus P4-induced transcriptional upregulation of PTGER2, results in a transition to a PKA-dominated, rather than AKT- dominated, signaling state.

Progestin-dependent expression of *PTGER2* has previously been reported in human cells (Brar et al., 1997). Dependence of decidualization upon progesterone has been found to be the result of cAMP-activating PGE2 receptor activation (i.e. PTGER2) in the rabbit (Fortier et al., 1987; Stadtmauer and Wagner, unpublished) and dog (Graubner et al., 2020). Both PTGER2 and PTGER4 are G-protein coupled receptors that signal through the Gαs subunit which activates adenylyl cyclase to produce cAMP. However, PTGER4 activates two signal transduction pathways, protein kinase A (PKA), which promotes decidualization, and phosphoinositide 3-kinase/protein kinase B (PI3K/AKT), which opposes it (Rundhaug et al., 2011; Gellersen and Brosens, 2014). Of the two kinases, PTGER4 favors AKT (Fujino et al., 2005), whereas PTGER2 is specific for PKA (Fujino et al., 2002). Decidualization is therefore unlikely to be inducible by PTGER4 signaling alone, given the demonstrated necessity of PKA signaling. Instead, it is likely that upon induction of PTGER2 and desensitiziation of PTGER4, preferential signaling through PTGER2 favors a PKA-dominated intracellular state, and thus promotes decidualization.

PTGER2 has been characterized as the least expressed of the four paralogous PGE2 receptors (Ricciotti and Fitzgerald, 2011), suggesting that its physiological role is more specialized. In pulmonary fibroblasts, prostaglandin E2 signaling through PTGER2 inhibits PI3K/AKT signaling, the myofibroblast phenotype and fibrosis (Mukherjee et al., 2019). Through PGE2-PTGER2 signaling, the same antagonistic mechanism may channel decidualizing endometrial stromal cells away from a more myofibroblast-like ESF state. In our single-cell expression data, *PTGER2* was associated with the mature decidual end of the decidualization curve, and had little to no intersection with markers of the acESF state (Figure 4). Phosphorylated (activated) AKT has been shown to decrease in cells decidualizing under the influence of progesterone, but phospho-AKT is elevated in the absence of progesterone or in the presence of progesterone plus a PKA inhibitor (Yoshino et al., 2003). This is consistent with a progesterone-dependent mechanism for shifting the balance from AKT-dominated to PKA-dominated state such as the one proposed here.

### Cellular Heterogeneity and Prostaglandin E2-Driven Decidualization

Decidual cell heterogeneity is the hypothesis that decidualization gives rise to multiple cell types, or multiple states of the same cell type. Different cell populations may coexist at the same time and fulfill distinct biological functions, in contrast to (but compatible with) the notion of temporal phases, including an early pro-inflammatory and later mature decidual state (Salker et al., 2012; Gellersen and Brosens, 2014; Rytkönen et al., 2019).

Single-cell RNA sequencing studies of the peri-implantation fetal-maternal interface (Vento- Tormo et al., 2018; Suryawanshi et al., 2018) support the model of decidual cell heterogeneity, with each partitioning DSC-like cells into three distinct transcriptional states. In the analysis by Vento- Tormo and colleagues (2018), these included a cluster lacking *PRL* and *IGFBP1* expression but expressing *ACTA2* and *TAGLN*, markers of the acESF cells identified in our study particularly prevalent at 2 days, as well as two mature decidual cell population characterized by expression of standard markers *PRL* and *IGFBP1*, although to a higher degree in one cluster with the other expressing markers such as *LEFTY2* and *IL15*. Notably, *LEFTY2* was one of the genes we found more highly elevated in PGE2+MPA treatment than cAMP+MPA, and the majority of the DSC cluster identified at 6 days was *LEFTY2^+^* and *IL15^+^*, with an offshoot more highly expressing *PRL* and *SST.* The authors implied a possible identification of the *LEFTY2^+^* cluster as snDSC although their justification was unclear

(Vento-Tormo et al., 2018). Our *LEFTY2^+^* cells from in vitro PGE2+MPA treatment lack expression of the putative senescence markers, so it is unlikely that the reason we find a single cluster of DSC is because *all* DSC are snDSC. The difference between these two DSC populations was not deemed great enough to warrant unique cluster assignment (appearing at k ≥ 5 clusters, Figure S4), but nevertheless this intriguing if weak pattern will need to be pursued by extended sampling. In the analysis by Suryawanshi and colleagues, the three subtypes of DSC were analyzed explicitly as a developmental trajectory, with one being an intermediate more ESF-like starting point, an *ACTA2^+^* and *TAGLN^+^* state, and a canonical decidual state expressing *PRL* and *IGFBP1.* Overall, comparison between the present study and the two in vivo analyses suggests that the most robust pattern is the identification of canonical *PRL^+^ IGFBP1^+^* DSC and an *ACTA2^+^ TAGLN^+^* cell we deem activated ESF.

The leading form of decidual cell heterogeneity discovered in vitro is what has been labeled as the senescent decidual cell state, proposed to have a physiological role in decidual development (Brighton et al., 2017) as well as a pathological role in recurrent pregnancy loss (Lucas et al., 2020). According to this theory, development of decidual stromal cells along a senescence pathway promotes tissue turnover and makes room for the implantation of the blastocyst, and is aided by the active phagocytic and cytolytic activities of uterine natural killer cells. Single-cell transcriptomic studies of in vitro decidualized human endometrial stromal fibroblasts revealed a trajectory splitting event at 4 days of decidualization after which “mature” and “senescent” decidual cells diverge (Lucas et al., 2020), but used the cAMP+MPA decidualization regime.

Our PGE2-mediated decidualization experiment focussed on 2- and 6-day time points to bracket this splitting event at day 4 reported by Lucas and collabortors (2020). Nevertheless, we did not identify a unique cluster cells co-expressing decidual and senescent markers after six days of PGE2+MPA treatment, although some of the cells we deem activated ESF did express a suite of genes associated with so-called snDSC in the absence of decidual markers. The lack of discretely identifiable snDSC after PGE2+MPA treatment could be interpreted as indicating that their induction under cAMP+MPA is an artifact of the stimulation with extracellular cAMP. Alternatively, the lack of senescent cells could be an artifact of the PGE2+MPA treatment. However, since our experiments show that decidualization by PGE2+MPA treatment is acting through the PTGER2 → cAMP → PKA pathway, we are inclined to believe that treatment with extracellular cAMP may overload the cell with cAMP, leading to cell senescence.

The senescence DSC model of Lucas et al. makes several assertions with which we agree. First, a group of genes including *ABI3BP*, *ADAMTS5*, *DIO2*, *CLU*, and *CEMIP* show correlated expression with *SOX4* or *STAT1.* Our data support this pattern, which is stronger at 6 days relative to 2 days.

Second, with respect to the finding that cells with this gene expression pattern increase after 4 days, our PGE2+MPA data also show expression of several of these genes at 2 and 6 days, although to a lesser degree than seen after cAMP+MPA treatment and not co-expressed with decidual markers. As a consequence, the finding that these cells are the product of a branching event or new endpoint that emerges after 4 days was not supported in our study.

The senescent and mature decidual cells can also be framed as a dichotomy between progesterone-resistant and progesterone-susceptible cells (Lucas et al., 2020), due to snDSC transcription factors being negatively regulated by progesterone (e.g. SOX4: Cloke et al., 2008). Since the upstream components of the decidualization pathway (such as PTGER2) are demonstrated to be regulated by progesterone signaling, the production of cells entering this state under the cAMP+MPA method could be increased as a consequence of exogenous cAMP bypassing this layer of regulation.

The findings from our study are compatible with the following model (Figure 7): Endometrial stromal fibroblasts decidualizing in response to PGE2 go through a pro-inflammatory phase in which they may assume an activated state similar to the myofibroblast-like transcriptomic profile identified in vivo. At 2 days, most of the cells are either in this state or in transition to DSC, whereas by 6 days many have adopted a more mature, canonical DSC state although some remain in the activated ESF state. Activated ESF are enriched for genes in the protein kinase B pathway and cell adhesion, whereas cells at or approaching the DSC state express *PTGER2*, the receptor through which PGE2 activates cAMP and protein kinase A signaling, even by day 2. At 6 days, activated ESF exist along a continuum where some have taken on a state where they express the putative senescence markers, PI3K/AKT- associated genes, and genes related to control of cell survival and adhesion (snESF). If 8-Br-cAMP is used as the deciduogenic stimulus, and if the snESF phenotype is indeed indicative of progesterone resistance, the production of snDSC under 8-Br-cAMP treatment but not PGE2+MPA can be explained by the bypassing of the surface receptor level of signal transduction upstream of cAMP/PKA. As PGE2 receptor expression, such as the EP2 switch we identify, seem to be influenced by progesterone, direct cAMP addition allows for the expression of the normally progesterone-dependent decidual program with the senescence-like program, resulting in snDSC.

**Figure 7.**
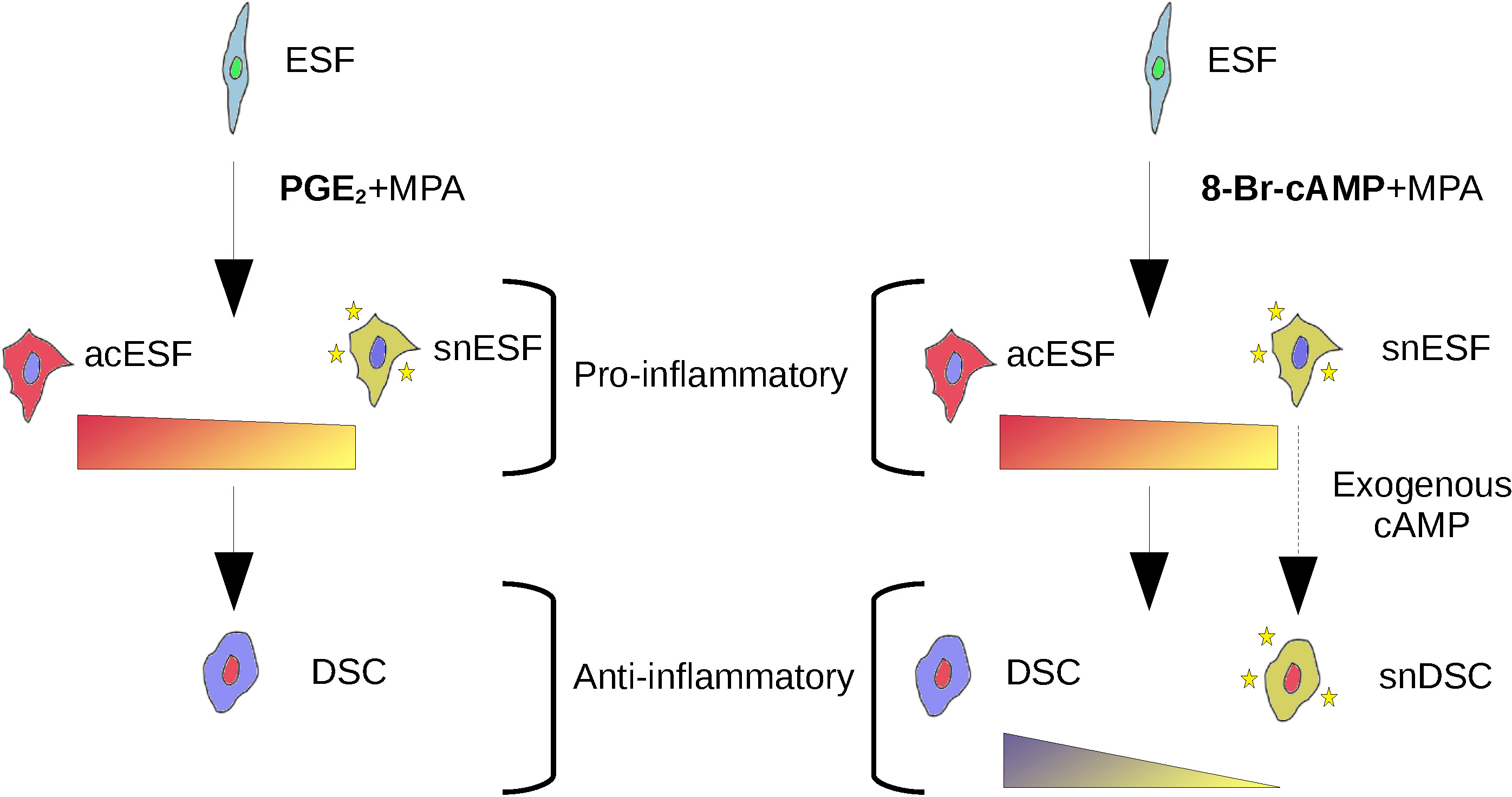
Conceptual diagram of cell states and types produced during in vitro decidualization with the PGE2+MPA (left) or 8-Br-cAMP+MPA (right) protocols. Three categorical cell states of (proliferative) ESF, activated ESF (acESF), and decidual stromal cells (DSC). Activated ESF exist along a continuum, one end of which (snESF) expresses a “senescence” gene program associated with stress-susceptibility/progesterone-resistance; under 8-Br-cAMP+MPA treatment these can become senescent DSC (snDSC) whereas under the PGE2+MPA model the single inferred endpoint is the canonical DSC.

Our results are consistent with the notion of temporal phases to decidualization, where decidual cells must progress from a proliferative fibroblast-like state to a pro-inflammatory and ultimately an anti-inflammatory state. Such a transition may require a distinct cellular program of resolution (Serhan and Savill, 2005). Notably, resolution of inflammation involves cell turnover through auto-opsoniaztion with cell surface molecules such as thrombospondin 1 (Serhan and Savill, 2005) which promotes immune cell recognition and phagocytosis of inflammatory cells. Indeed, *THBS1* was a top marker of the activated ESF cell state, along with the thrombospondin motif-containing *ADAMTS5* enriched in snESF. Since this phenomenon can occur in many cell types, its evolution predates the decidual stromal cell and thus it may be expected that development towards an anti-inflammatory state would activate programs for, if not direct cell death, then immune-assisted cell death, a cornerstone of the senescence model (Brighton et al., 2017). Virtually none of the recognized products of the acute senescence-associated secreteory phenotype (Coppé et al., 2010) are expressed in our data.

In vitro decidualization is inevitably removed from in vivo biology. As it has become evident that the decidual stromal cell type takes several forms depending upon developmental time and situation (e.g. blastocyst rejection or acceptance), when choosing an in vitro model system it is important to note which of these states it produces. Our PGE2+MPA in vitro decidualization appears to generate mature decidual stromal cells as well as a subset of activated ESF similar to the myofibroblast-like activated ESF identified in in vivo studies (Suryawanshi et al., 2018; Vento-Tormo et al., 2018). On the other hand, PGE2+MPA decidualization appears to less readily generate senescent DSC. Should senescent DSC be a biologically relevant entity, the identification and use of natural ligands for this specific cell state could be an improvement to their generation in response to exogenous cAMP. Method choice would then depend upon the endometrial state one intends to simulate in vitro.

Importantly, our failure to reliably generate snDSC using PGE2+MPA does not contradict the possibility that stromal cell senescence takes place in vivo and can contribute to recurring early pregnancy loss (Lucas et al., 2020).

### The Evolution of Decidual Cell Differentiation

According to the sister cell type model (Arendt, 2008), novel cell types come into existence through an evolutionary bifurcation event which results in the production of two distinct cell types to take the place of the ancestral cell type – hence, sister cell types. Within this framework, the neo-ESF cell of placental mammals is the sister cell type to the decidual stromal cell (Kin *et al.,* 2015). In this case one sister cell type, the neo-ESF, also gives rise to the other, the DSC, during ontogeny. This developmental linkage is itself a derived state: exposure of opossum paleo-ESF to deciduogenic conditions – either cAMP+MPA or PGE2+MPA – results in cellular stress rather than decidualization (Erkenbrack *et al.,* 2018). Most notably, the same study also provides data that shows that MPA does not induce the expression of PTGER2 in the opossum ESF, suggesting that the progesterone responsiveness of the *PTGER2* gene is a key innovation in the evolution of the neo-ESF/DSC sister cell type pair. These findings suggest that it was only upon the evolution of progesterone-dependent regulation of PTGER2 that the two natural ligands for decidualization, PGE2 and progesterone, become functionally linked. This linkage may be a way in which the progesterone channels the inflammation induced PGE2 expression into a signal for decidual cell type differentiation, rather than stress and senescence.

## Acknowledgments

DJS is thankful to E. Erkenbrack for guidance in cell culture procedures and experimental design. The analytical component of this study benefited greatly due to computational guidance from, and software made available by, D. Burkhardt and S. Gigante of the Krishnaswamy lab as well as A. Chavan and A. Dighe of the Wagner lab. The authors recognize the support of the National Cancer Institute (U54-CA209992) and John Templeton Foundation (#61329). DJS is supported by the NIH Predoctoral Training Program in Genetics (T32 GM 007499). The opinions expressed in this paper are those of the authors and do not represent the stance of the funding institutions.

## Table

**Table S1.**
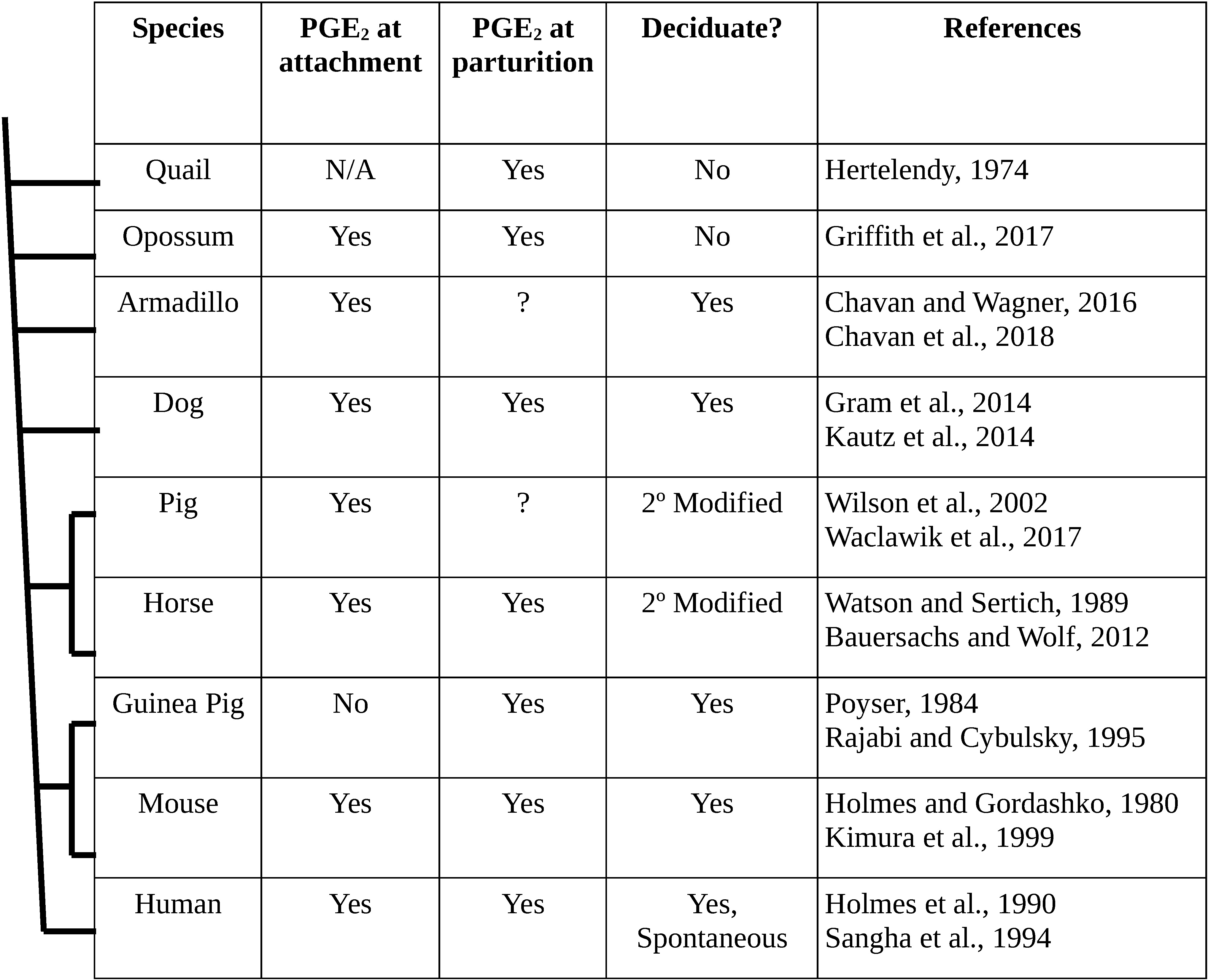
Distribution of prostaglandin E2 in pregnancy (attachment and parturition) in selected amniotes, with phylogenetic relationships sketched on the left. Cursory ancestral state inference suggests that PGE2 involvement in attachment predated the evolution of the decidua, consistent with PGE2 being a natural ligand for decidualization.

| Species | PGE<sub>2</sub> at attachment | PGE<sub>2</sub> at parturition | Deciduate? | References |
| --- | --- | --- | --- | --- |
| Quail | N/A | Yes | No | Hertelendy, 1974 |
| Opossum | Yes | Yes | No | Griffith et al., 2017 |
| Armadillo | Yes | ? | Yes | Chavan and Wagner, 2016<br>Chavan et al., 2018 |
| Dog | Yes | Yes | Yes | Gram et al., 2014<br>Kautz et al., 2014 |
| Pig | Yes | ? | 2° Modified | Wilson et al., 2002<br>Waclawik et al., 2017 |
| Horse | Yes | Yes | 2° Modified | Watson and Sertich, 1989<br>Bauersachs and Wolf, 2012 |
| Guinea Pig | No | Yes | Yes | Poyser, 1984<br>Rajabi and Cybulsky, 1995 |
| Mouse | Yes | Yes | Yes | Holmes and Gordashko, 1980<br>Kimura et al., 1999 |
| Human | Yes | Yes | Yes,<br>Spontaneous | Holmes et al., 1990<br>Sangha et al., 1994 |

**Table S2.** Centers of standardized distributions of expression value scores (Z-score derived from square root TPM) for each gene cluster as determined by k-means clustering. For the purposes of interpretation, these values values can be binned into “low/off” (z < -0.33), “medium” (-0.33 < z < +0.33), and “high” (z > +0.33) levels of expression.

| Cluster | Base | cAMP+MPA | PGE<sub>2</sub>+MPA |
| --- | --- | --- | --- |
| 1 | +0.85 (H) | -1.06 (L) | +0.21 (M) |
| 2 | +1.12 (H) | -0.75 (L) | -0.37 (L) |
| 3 | -0.87 (L) | +1.05 (H) | -0.17 (M) |
| 4 | -1.09 (L) | +0.49 (H) | 0.60 (H) |
| 5 | +1.01 (L) | -0.10 (M) | -0.90 (L) |

**Table S3.** Expression levels of select genes functionally related to decidual cell identity or phenotype, all falling into Cluster 4 in k-means clustering analysis.

| Gene Name | TPM<br>(Base) | TPM<br>(cAMP) | TPM<br>(PGE<sub>2</sub>) | FDR (Base<br>vs. PGE<sub>2</sub>) | FDR (Base<br>vs. cAMP) | FDR<br>(PGE<sub>2</sub> vs.<br>cAMP) |
| --- | --- | --- | --- | --- | --- | --- |
| *FOS* | 4.1 | 6.5 | 7.1 | 1.2E-10 | 8.4E-04 | 5.3E-03 |
| *FOXO1* | 1.9 | 77.2 | 43.4 | 0.0E+00 | 0.0E+00 | 6.0E-26 |
| *HAND2* | 33.4 | 157.3 | 114.4 | 4.2E-181 | 1.2E-252 | 1.2E-06 |
| *HIF1A* | 174.6 | 365.1 | 364.0 | 2.8E-61 | 2.5E-48 | 1.1E-01 |
| *KLF9* | 5.6 | 19.0 | 23.7 | 1.2E-160 | 3.1E-94 | 1.1E-10 |
| *NR4A2* | 0.7 | 13.0 | 9.5 | 1.5E-59 | 1.4E-65 | 5.1E-01 |
| *OSR2* | 18.8 | 44.7 | 38.9 | 1.9E-27 | 3.6E-31 | 6.5E-01 |
| *ZBTB16* | 0.1 | 4.4 | 4.8 | 1.4E-36 | 9.5E-33 | 5.9E-01 |
| *ZHX2* | 4.4 | 10.5 | 8.9 | 4.6E-19 | 2.2E-18 | 8.8E-01 |
| *ZNF331* | 7.0 | 14.5 | 12.0 | 2.2E-22 | 1.9E-23 | 9.4E-01 |
| *BMP2* | 0.5 | 6.5 | 4.5 | 8.2E-34 | 2.2E-43 | 1.4E-01 |
| *IGFBP1* | 0.0 | 4.8 | 7.5 | 3.3E-33 | 2.9E-25 | 5.2E-02 |
| *LEFTY2* | 0.0 | 14.5 | 40.2 | 4.7E-85 | 1.4E-56 | 1.4E-08 |
| *PRL* | 0.2 | 13.3 | 9.7 | 1.5E-26 | 2.9E-31 | 4.7E-01 |
| *SST* | 0.1 | 8.3 | 221.1 | 3.9E-195 | 2.4E-20 | 1.6E-125 |

**Table S4.** Expression levels of select genes proposed to be markers of the senescent decidual cell state (Lucas et al., 2020), all falling into Cluster 3 in k-means clustering analysis.

| Gene Name | TPM<br>(Base) | TPM<br>(cAMP) | TPM<br>(PGE<sub>2</sub>) | FDR (Base<br>vs. PGE<sub>2</sub>) | FDR (Base<br>vs. cAMP) | FDR<br>(PGE<sub>2</sub> vs.<br>cAMP) |
| --- | --- | --- | --- | --- | --- | --- |
| *ABI3BP* | 37.6 | 469.7 | 211.3 | 6.4E-140 | 2.1E-248 | 1.1E-22 |
| *ADAMTS5* | 140.3 | 216.8 | 122.2 | 5.9E-02 | 3.8E-34 | 5.6E-46 |
| *CEMIP* | 6.9 | 40.7 | 21.0 | 4.2E-08 | 2.5E-18 | 2.3E-03 |
| *CLU* | 154.8 | 436.1 | 136.8 | 6.1E-01 | 8.4E-28 | 2.0E-30 |
| *DIO2* | 11.6 | 25.8 | 20.0 | 8.2E-04 | 9.4E-08 | 1.3E-01 |
| *IGF2* | 14.0 | 306.9 | 89.0 | 2.5E-71 | 1.4E-159 | 4.4E-28 |
| *LUM* | 81.0 | 377.6 | 250.4 | 8.7E-94 | 5.9E-144 | 1.3E-06 |
| *NPC2* | 692.7 | 1962.9 | 1369.2 | 3.6E-35 | 4.9E-63 | 5.2E-05 |
| *SOX4* | 111.8 | 176.3 | 99.0 | 1.8E-01 | 7.9E-22 | 1.6E-28 |

**Table S5.** Expression levels of genes falling into Cluster 5, with reduced expression after PGE2+MPA treatment compared to base or cAMP+MPA treatment.

| Gene Name | TPM<br>(Base) | TPM<br>(cAMP) | TPM<br>(PGE<sub>2</sub>) | FDR (Base<br>vs. PGE<sub>2</sub>) | FDR (Base<br>vs. cAMP) | FDR<br>(PGE<sub>2</sub> vs.<br>cAMP) |
| --- | --- | --- | --- | --- | --- | --- |
| *NRG1* | 48.7 | 62.4 | 20.9 | 5.0E-47 | 4.8E-19 | 5.3E-122 |
| *GFRA1* | 33.3 | 25.0 | 13.2 | 1.3E-117 | 2.6E-13 | 3.3E-53 |
| *PTN* | 189.4 | 162.4 | 63.9 | 5.4E-76 | 4.1E-04 | 1.9E-48 |
| *TLE4* | 223.2 | 135.3 | 71.6 | 5.2E-135 | 2.1E-30 | 3.8E-39 |
| *HAS2* | 8.9 | 11.3 | 4.0 | 3.0E-20 | 1.3E-02 | 5.6E-32 |
| *MEX3A* | 12.4 | 13.8 | 6.4 | 1.4E-17 | 3.4E-02 | 5.9E-27 |
| *PCDH9* | 14.7 | 10.3 | 5.1 | 1.2E-79 | 3.6E-16 | 4.3E-25 |
| *GBE1* | 78.3 | 74.6 | 44.2 | 1.1E-35 | 1.4E-02 | 5.2E-22 |
| *PAG1* | 13.6 | 11.4 | 8.2 | 4.4E-30 | 1.1E-01 | 3.9E-21 |
| *LRRC17* | 52.7 | 21.0 | 7.6 | 6.2E-92 | 2.3E-30 | 4.4E-21 |
| *AC055839.2* | 84.6 | 65.4 | 33.0 | 7.1E-49 | 3.3E-07 | 3.2E-20 |
| *TRIM5* | 25.6 | 31.6 | 16.4 | 6.9E-09 | 2.3E-03 | 1.6E-19 |
| *ZNF804A* | 16.8 | 9.5 | 4.5 | 7.6E-84 | 7.0E-26 | 5.7E-19 |
| *DCHS1* | 11.5 | 7.5 | 3.5 | 3.9E-58 | 4.8E-12 | 9.2E-19 |
| *SPP1* | 21.4 | 27.1 | 7.7 | 4.6E-12 | 1.7E-01 | 7.1E-17 |

---

**Figure S1.**
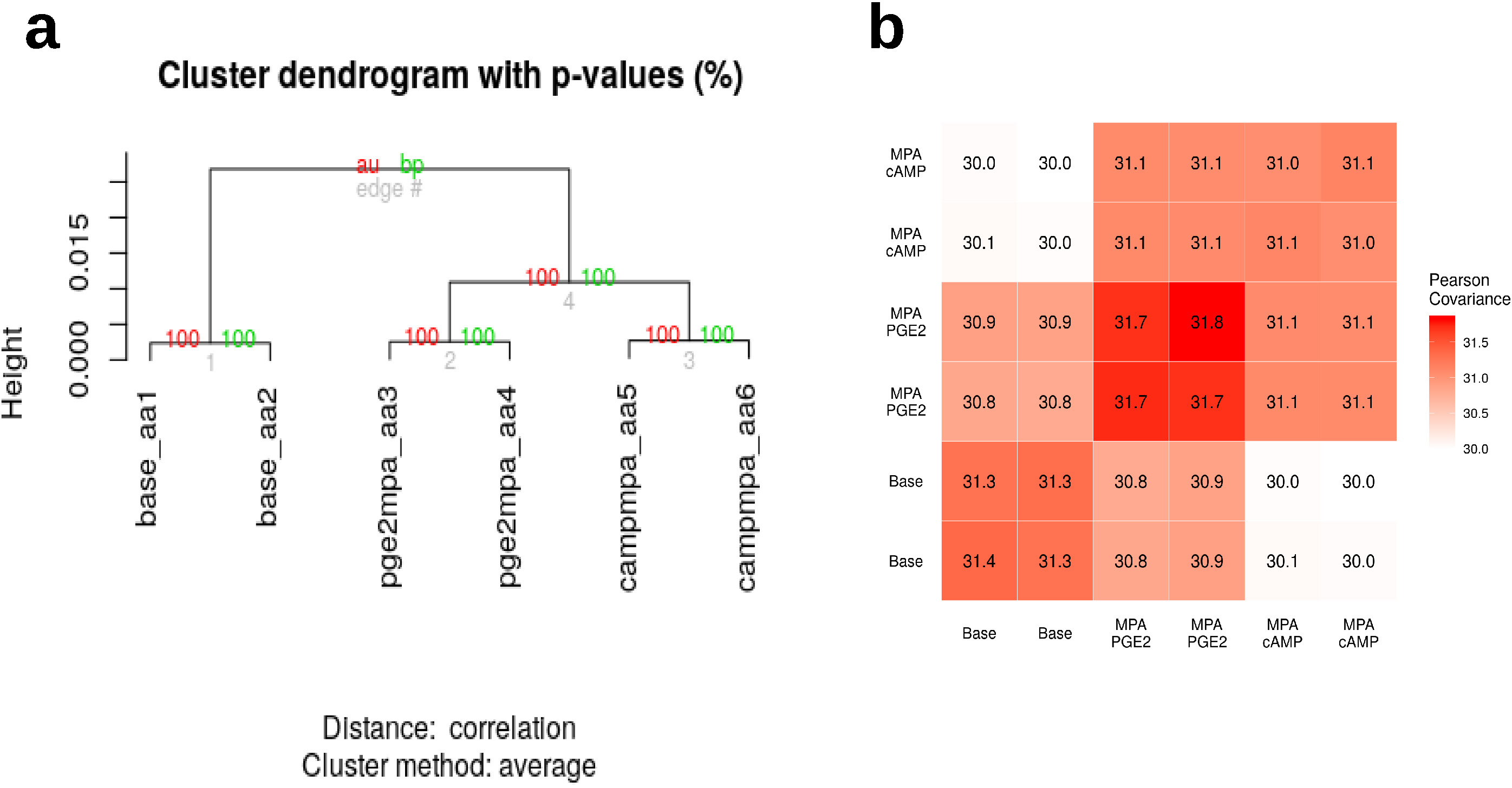
Bulk RNA sequencing analysis. **a.** Clustering dendrogram of all genes for 3 treatment conditions ran in duplicate. **b.** Covariance matrix between protein-coding genes of the 6 bulk transcriptomes generated.

**Figure S2.**
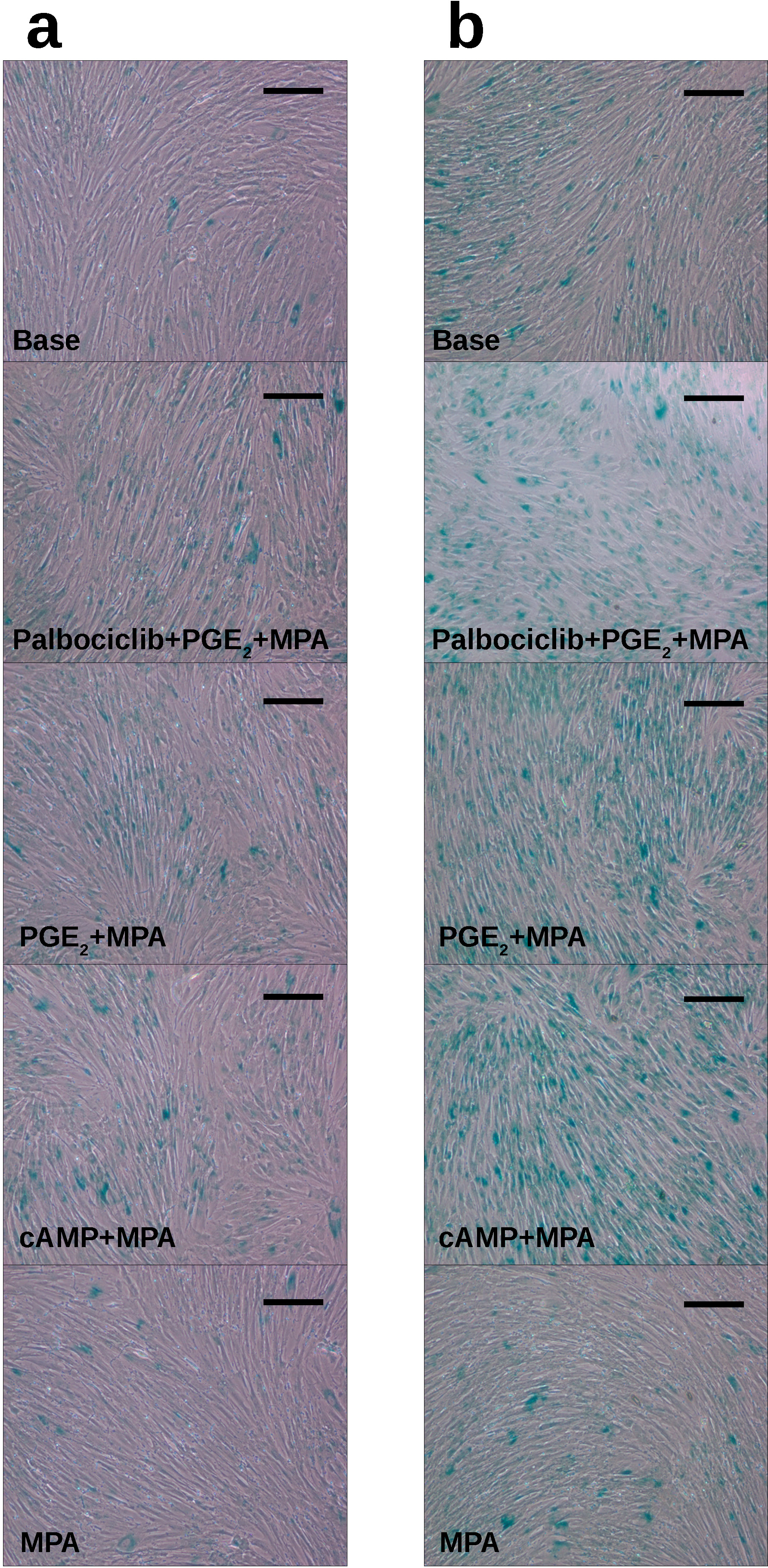
Senescence-associated β-galactosidase staining. **a.** Representative images from cells treated for 3 days (1 of 2 replicates ran). **b.** Representative images from cells treated for 6 days. Scale bar = 20 μm.

**Figure S3.**
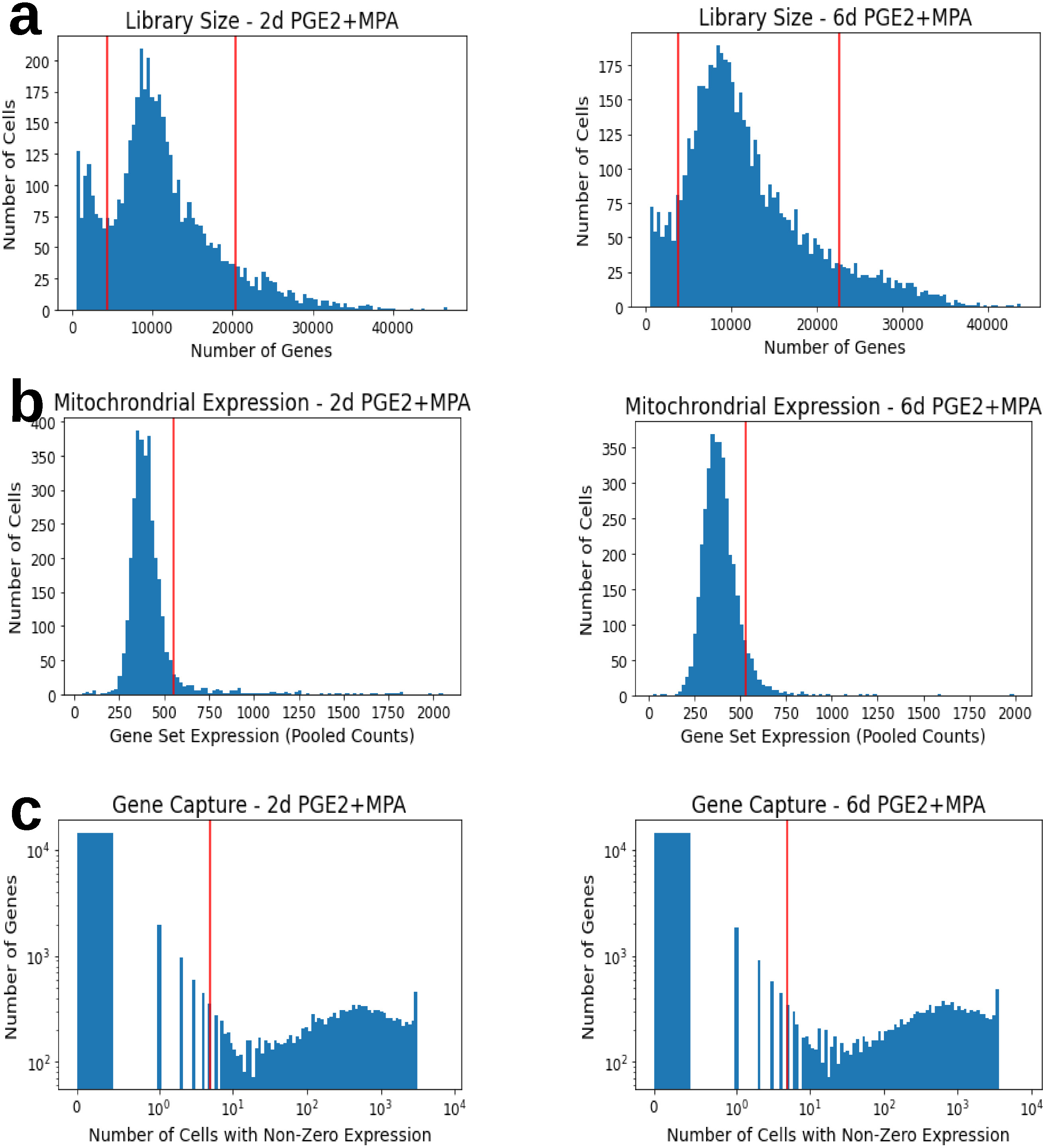
Quality control metrics and filtering cutoffs for single-cell RNA sequencing analysis. **a.** Library size cutoffs were chosen for each sample (2- day,left: 16%,90%; 6-day, right: 9%,90%) by a visual attempt to isolate a single symmetrical distribution by removal of a low-size peak and removal of a high library size tail. **b.** Mitochondrial gene expression was cut off at the 93^rd^ percentile of both samples in combination. **c.** Genes expressed in fewer than 5 cells in total from both samples were excluded.

**Figure S4.**
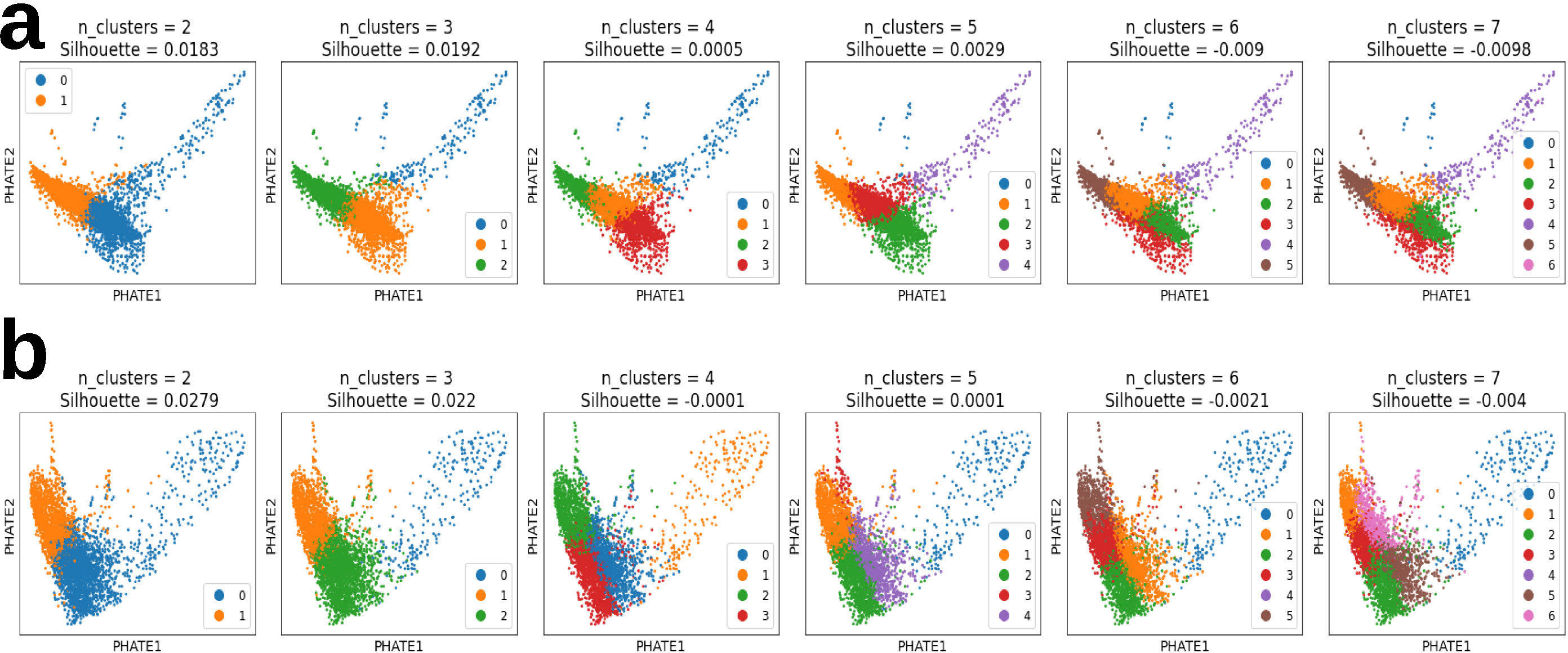
K-means clustering of transcriptomes at 2 days (**a**) and 6 days (**b**) was performed with k parameters from 2 to 7 clusters. Silhouette = silhouette score.

**Figure S5.**
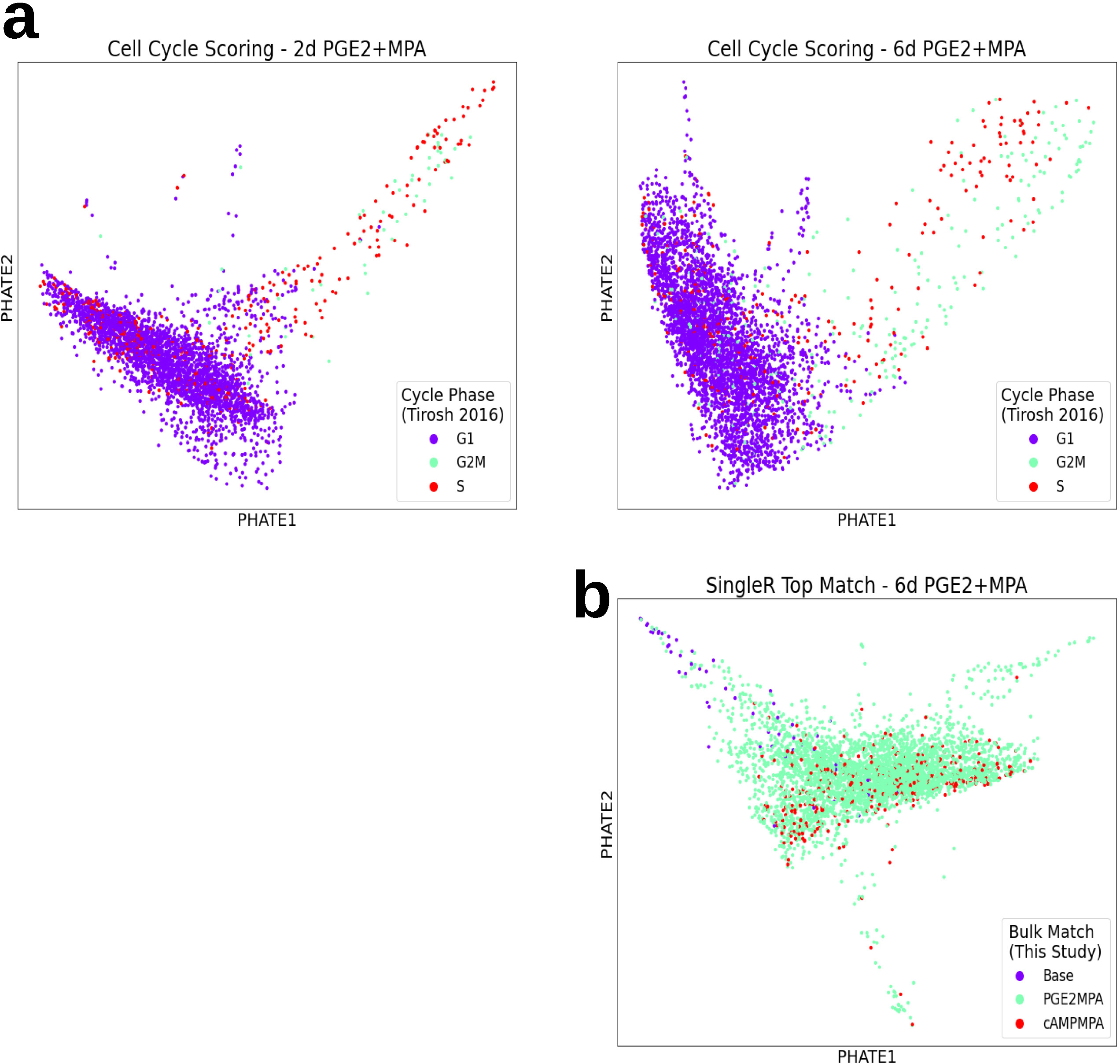
Cell cycle inference. **a.** Scoring of most likely cell cycle phase, based upon marker genes identified by Tirosh and colleagues (2016). **b.** Regression out of cell cycle-associated genes did not remove the sparse tail of ESF-like cells, so it was not performed in the primary analysis.

**Figure S6.**
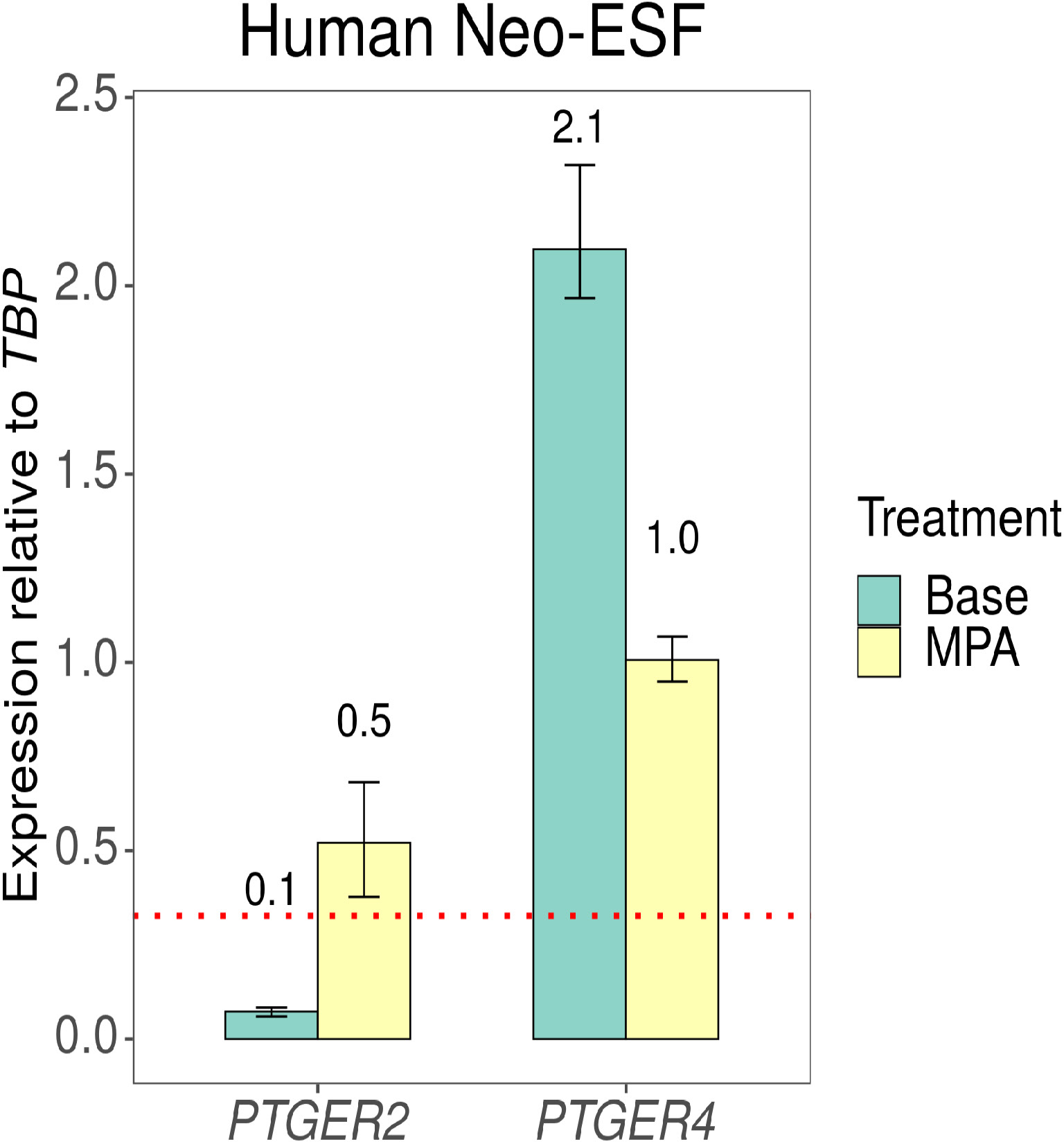
Additional qPCR quantification of *PTGER2* and *PTGER4* endometrial stromal fibroblasts after 6 days of daily treatment with media ± MPA. *PTGER2* increased 7.14-fold in response to MPA, passing from “off” to “on”, whereas *PTGER2* was reduced to 48% of its original value. Red line (y-intercept = 0.327) shows approximate threshold for being actively transcribed, based on an average TPM value of 9.17 for *TBP* (this study) and cutoff of 3 TPM. Experimental n = 2 for all conditions and technical n = 3.

